# Molecular basis of sulfolactate synthesis by sulfolactaldehyde dehydrogenase from *Rhizobium leguminosarum*

**DOI:** 10.1101/2023.03.13.532361

**Authors:** Jinling Li, Mahima Sharma, Richard Meek, Amani Alhifthi, Zachary Armstrong, Niccolay Madiedo Soler, Mihwa Lee, Ethan D. Goddard-Borger, James N. Blaza, Gideon J. Davies, Spencer J. Williams

## Abstract

Sulfolactate (SL) is a short-chain organosulfonate that is an important reservoir of sulfur in the biosphere. SL is produced by oxidation of sulfolactaldehyde (SLA), which in turn derives from sulfoglycolysis of the sulfosugar sulfoquinovose, or through oxidation of 2,3-dihydroxypropanesulfonate. Oxidation of SLA is catalyzed by SLA dehydrogenases belonging to the aldehyde dehydrogenase superfamily. We report that SLA dehydrogenase *Rl*GabD from the sulfoglycolytic bacterium *Rhizobium leguminsarum* SRDI565 can use both NAD^+^ and NADP^+^ as cofactor to oxidize SLA, and indicatively operates through a rapid equilibrium ordered mechanism. We report the cryo-EM structure of *Rl*GabD bound to NADH, revealing a tetrameric quaternary structure and supporting proposal of organosulfonate binding residues in the active site, and a catalytic mechanism. Sequence based homology searches identified SLA dehydrogenase homologs in a range of putative sulfoglycolytic gene clusters in bacteria predominantly from the phyla Actinobacteria, Firmicutes, and Proteobacteria. This work provides a structural and biochemical view of SLA dehydrogenases to complement our knowledge of SLA reductases, and provide detailed insights into a critical step in the organosulfur cycle.

**Graphical abstract:** Sulfolactate is an important species in the biogeochemical sulfur cycle. Herein we report the 3D cryo-EM structure and kinetics of its biosynthetic enzyme, sulfolactaldehyde dehydrogenase.

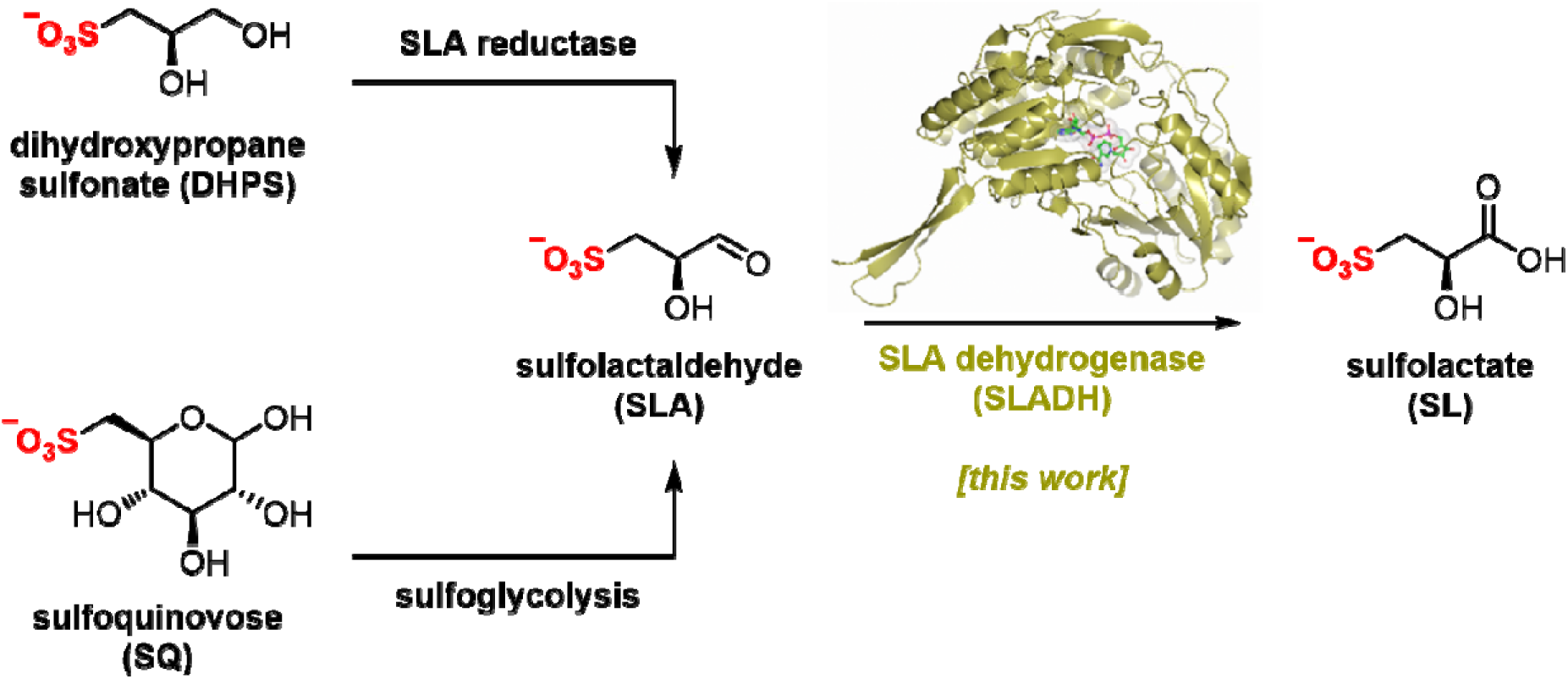

## Introduction

Sulfur is the tenth most common element by mass in the universe, the sixteenth most common in the Earth’s crust, and the sixth most abundant in seawater.^1^ It joins nitrogen, phosphorus, and potassium as the fourth macronutrient required for plants. The breakdown of C3-organosulfonates, primarily 2,3-dihydroxypropanesulfonate (DHPS) and sulfolactate (SL), allows the recycling of the element sulfur.^2^ DHPS and SL are produced from the reduction or oxidation, respectively, of sulfolactaldehyde (SLA).^3^ SLA in turn is produced in the pathways of sulfoglycolysis, through which the C6-organosulfonate sulfoquinovose (SQ) is catabolized (**Fig. 1a**).^4^ Alternatively, SLA may be produced by anaerobic DHPS degrading bacteria through the oxidation of DHPS (**Fig. 1b**).^5^ Reduction of SLA to DHPS is catalyzed by SLA reductase, an NADH-dependent enzyme, which has been biochemically and structurally characterized.^6, 7^ On the other hand oxidation of SLA to SL is poorly studied, with only basic evidence for the formation of product in coupled assays (*vide infra*).

**Fig. 1.**
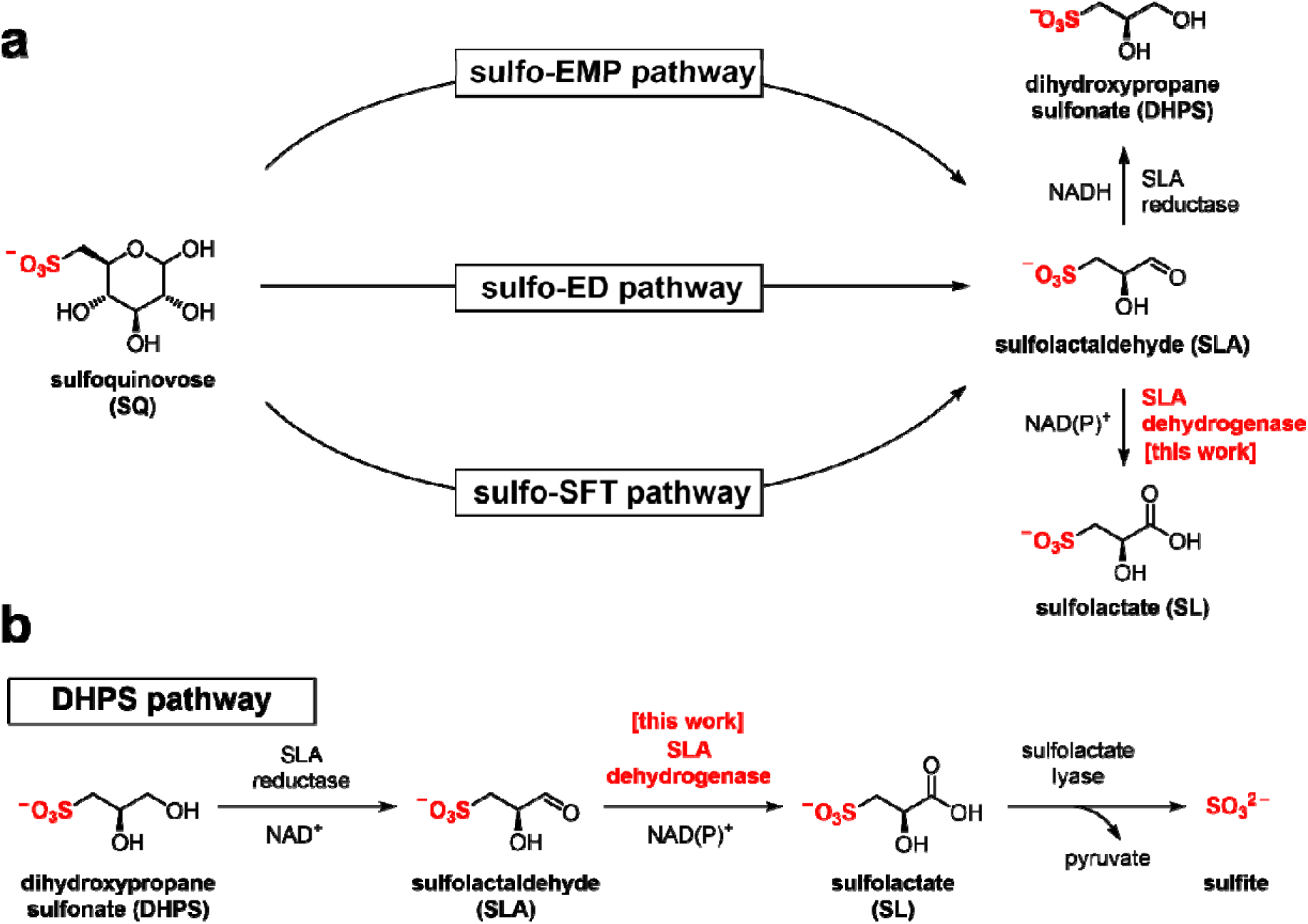
**(a)** Formation of SL and DHPS through the pathways of sulfoglycolysis from sulfoquinovose (SQ). **(b)** Formation and degradation of sulfolactate by catabolism of DHPS.

Three sulfoglycolytic pathways produce SLA by cleaving the 6-carbon chain of SQ into two C3 chains, namely the sulfoglycolytic Embden-Meyerhof-Parnas (sulfo-EMP/EMP2),^6, 8, 9^ Entner-Doudoroff (sulfo-ED)^10^ and sulfofructose transaldolase (sulfo-SFT) pathways (**Fig. 1a**).^11, 12^ These pathways generate dihydroxyacetone phosphate, pyruvate or fructose-6-phosphate (by transfer of a C3-glycerone moiety to glyceraldehyde-3-phosphate (GAP)), which are utilized by the host, and SLA, which is either reduced (to DHPS) or oxidized (to SL), and excreted. Examples of SL producing sulfoglycolytic organisms include: the sulfo-ED pathway (*Pseudomonas putida* SQ1,^10^ and *Rhizobium leguminosarum* bv. trifolii SRDI565^13^); the sulfo-EMP/EMP2 pathways (*Escherichia coli*,^6^ *Bacillus urumquiensis*,^7^ *Arthrobacter* spp.^9^); and the sulfo-SFT pathway (*Bacillus aryabhattai* SOS1,^11^ *Bacillus megaterium* DSM1804,^12^ and *Enterococcus gilvus*^11^). Gene clusters encoding these pathways are shown in **Fig. 2**. Excreted DHPS and SL are substrates for biomineralization bacteria. In the DHPS degradation pathway used by *Desulfovibrio* sp. strain DF1, DHPS is oxidized to SLA, and then SLA dehydrogenase SlaB oxidizes SLA to SL (**Fig. 1b**).^5^ SL is a substrate for SL lyase, which cleaves the C–S bond, producing pyruvate and sulfite.^14^ Other bacteria, such as *Roseovarius nubinhibens* and *Paracoccus pantotrophus*, utilize SL as a substrate for growth through the direct action of SL lyase.^5, 14, 15^

**Fig. 2.**
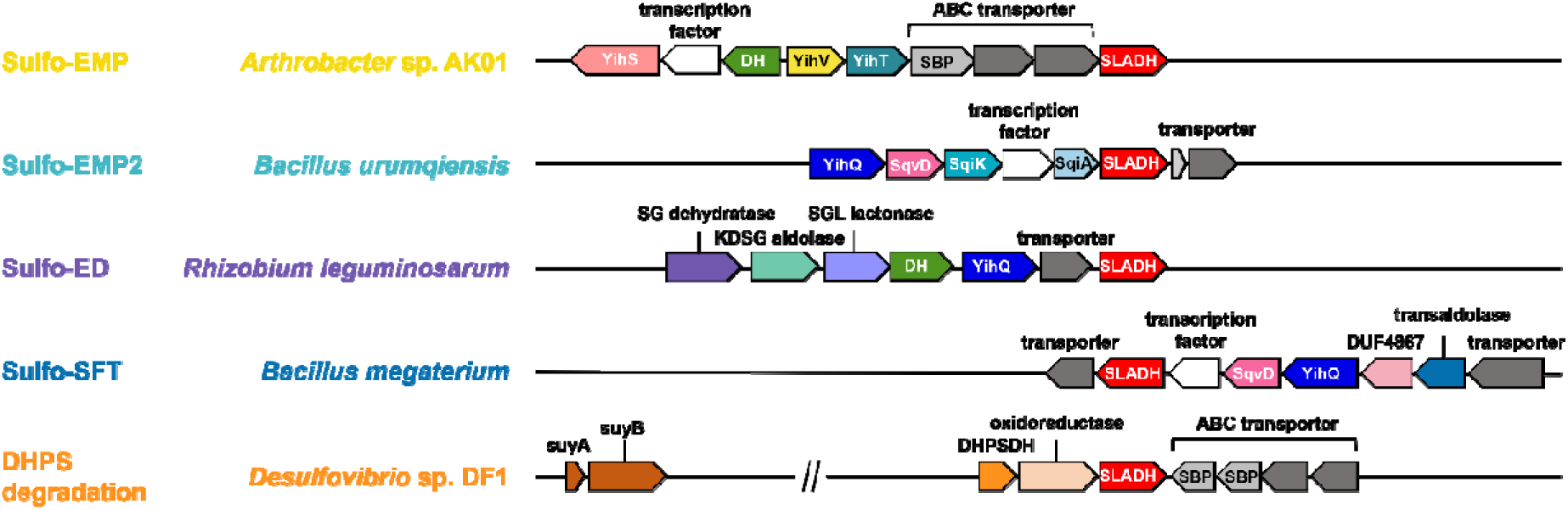
Proposed gene clusters of bacteria containing SLA dehydrogenase (SLADH) genes that degrade SQ (through sulfo-EMP, sulfo-EMP2, sulfo-ED, and sulfo-SFT pathways) and DHPS (through DHPS degradation pathway).

SLA dehydrogenases (annotated as GabD or SlaB) belong to the sequence-based protein family PF00171 within the Pfam database, which are members of the aldehyde dehydrogenase superfamily.^13^ Proteins of this superfamily oxidize the oxo group of aldehyde substrates to carboxylic acids, and use either NAD^+^ or NADP^+^ as hydride acceptors. Other activities within family PF00171 include succinate-semialdehyde dehydrogenase (SSADH),^14^ non-phosphorylating glyceraldehyde-3-phosphate dehydrogenase (GAPDH),^15^ and glutarate semialdehyde reductase.^16^ The potential cross-reactivity of SLA dehydrogenase with the structurally similar glycolytic intermediate glyceraldehyde-3-phosphate has not been reported.

Recombinant SLA reductase from *P. putida* SQ1 reduced SLA formed in situ in a coupled assay with both NAD^+^ and NADP^+^ cofactors,^7^ with a preference for the former, while SLA dehydrogenases from *B. aryabhattai* SOS1 (SftD),^11^ *B. megaterium* (SlaB)^10^ and *Desulfovibrio* sp. strain DF1 (SlaB)^5^ were described as NAD^+^ dependent, although it is unclear whether their ability to utilize NADP^+^ was assessed. In all cases accurate kinetic parameters have not been reported for any SLA dehydrogenase as SLA was not available in pure form. Recently, our group synthesized SLA from glycidol diethyl acetal using a chemical method,^16^ meaning that a comprehensive kinetic characterization of SLA dehydrogenase is now possible.

Here, we report the structure and reactivity of SLA dehydrogenase from *R. leguminasarum* SRDI565 (*Rl*GabD), which oxidizes SLA produced in a sulfo-ED pathway in this organism.^13^ We measure Michaelis-Menten kinetics and show its ability to use both NAD^+^/NADP^+^ as cofactors, its cross-reactivity to the structurally-related glycolytic metabolite GAP, and its sensitivity to inhibition by reduced NADH analogues. We determine its kinetic reaction order and provide evidence in support of an equilibrium ordered mechanism in the forward direction. We report the 3D structure of SLA dehydrogenase using CryoEM and define its quaternary structure and infer the SLA binding pocket, allowing proposal of a chemical mechanism of catalysis. Finally, we explore the sequence-based taxonomic distribution of SLA dehydrogenases across sulfoglycolytic and DHPS-degrading pathways using sequence similarity network analysis.

## Results and Discussion

### *Rl*GabD is an NAD(P)^+^-dependent SLA oxidase

The gene encoding GabD from *Rhizobium leguminosarum* (*Rl*GabD) was cloned, recombinantly expressed in *E. coli*, and purified to homogeneity. Reaction rates for the oxidation of SLA catalyzed by *Rl*GabD were measured using chemically-synthesized racemic D/L-SLA^16^ and monitoring reduction of NAD(P)^+^ to NAD(P)H using a UV/Vis spectrophotometer. Incubation of a solution of racemic SLA (1 mM) in Tris buffer with *Rl*GabD and excess NAD^+^ gave a progress curve that indicated complete reaction after 40 min; addition of more *Rl*GabD did not result in further conversion (**Fig. S1**). Based on the change in absorbance and the extinction coefficient for NAD^+^ we calculate that 47±2% of the SLA was consumed and conclude that *Rl*GabD is stereospecific for D-SLA. All subsequent analysis used the calculated D-SLA concentration (ie [SLA]/2).

Apparent Michaelis-Menten parameters were measured for D-SLA, NAD^+^ and NADP^+^ under pseudo first order conditions, in which one substrate was held at a constant concentration while that of the other was varied (**Fig. 3a-d**, **Table 1**). At 0.25 mM D-SLA, the pseudo first order parameters for NAD^+^ are: *k*_cat_^app^ = 17.7 s^−1^, *K*_M_^app^ = 0.081 mM and (*k*_cat_/*K*_M_)^app^ = 210 mM^−1^ s^−1^ and for NADP^+^: *k*_cat_^app^ = 4.1 s^−1^, *K*_M_^app^ = 0.017 mM and (*k*_cat_/*K*_M_)^app^ = 240 mM^−1^ s^−1^. Thus, while NADP^+^ has a lower *K*_M_^app^ value, the (*k*_cat_/*K*_M_)^app^ values of the two nucleotides are essentially identical. For variable D-SLA the Michaelis-Menten parameters at constant concentration (0.25 mM) of nucleotide were, NAD^+^: *k*_cat_^app^ = 17.8 s^−1^, *K*_M_^app^ = 0.13 mM and (*k*_cat_/*K*_M_)^app^ = 137 mM^−1^ s^−1^; and NADP^+^: *k*_cat_^app^ = 4.7 s^−1^, *K*_M_^app^ = 0.16 mM and (*k*_cat_/*K*_M_)^app^ = 30 mM^−1^ s^−1^. Comparison of (*k*_cat_/*K*_M_)^app^ reveal a modest preference for NAD^+^. Above 0.25 mM SLA, we observed substrate inhibition and so data were fit to rates measured at concentrations below this limit.

**Fig. 3.**
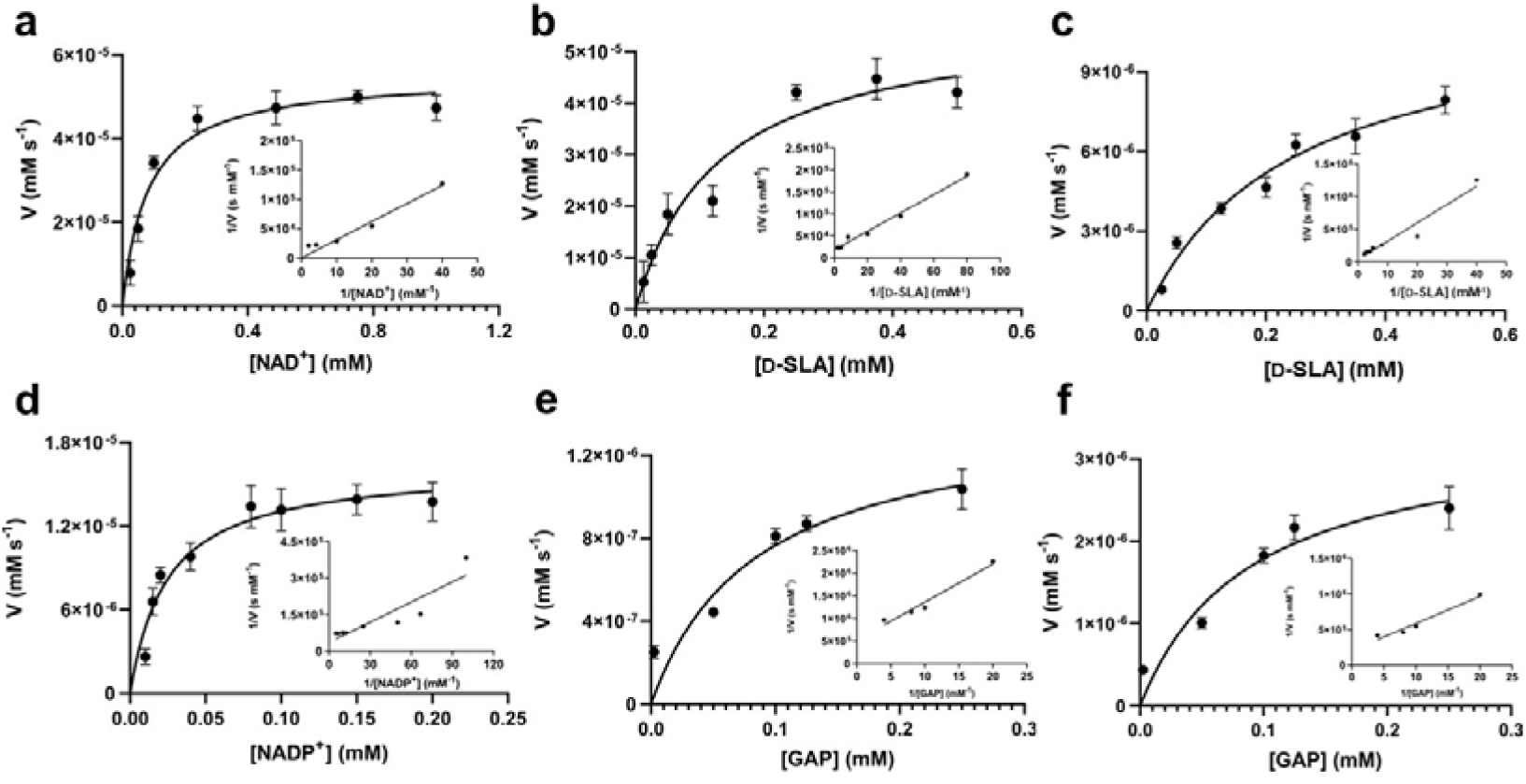
Michaelis-Menten kinetic analysis for SLA dehydrogenase using NAD^+^, D-SLA, NADP^+^ and GAP as substrates. **(a, b)** Michaelis-Menten and Lineweaver-Burk (inset) plots for *Rl*GabD under pseudo first-order conditions of [D-SLA] = 0.25 mM (for panel a) and [NAD^+^] = 0.25 mM (for panel b). **(c, d)** Michaelis-Menten and Lineweaver-Burk (inset) plots for *Rl*GabD under pseudo first-order conditions of [NADP^+^] = 0.25 mM (for panel c) and [D-SLA] = 0.25 mM (for panel d). **(e, f)** Michaelis-Menten and Lineweaver-Burk (inset) plots for oxidation of GAP by *Rl*GabD under pseudo first-order conditions of [NAD^+^] = 0.25 mM (for panel e) and [NADP^+^] = 0.25 mM (for panel f).

**Table 1.**
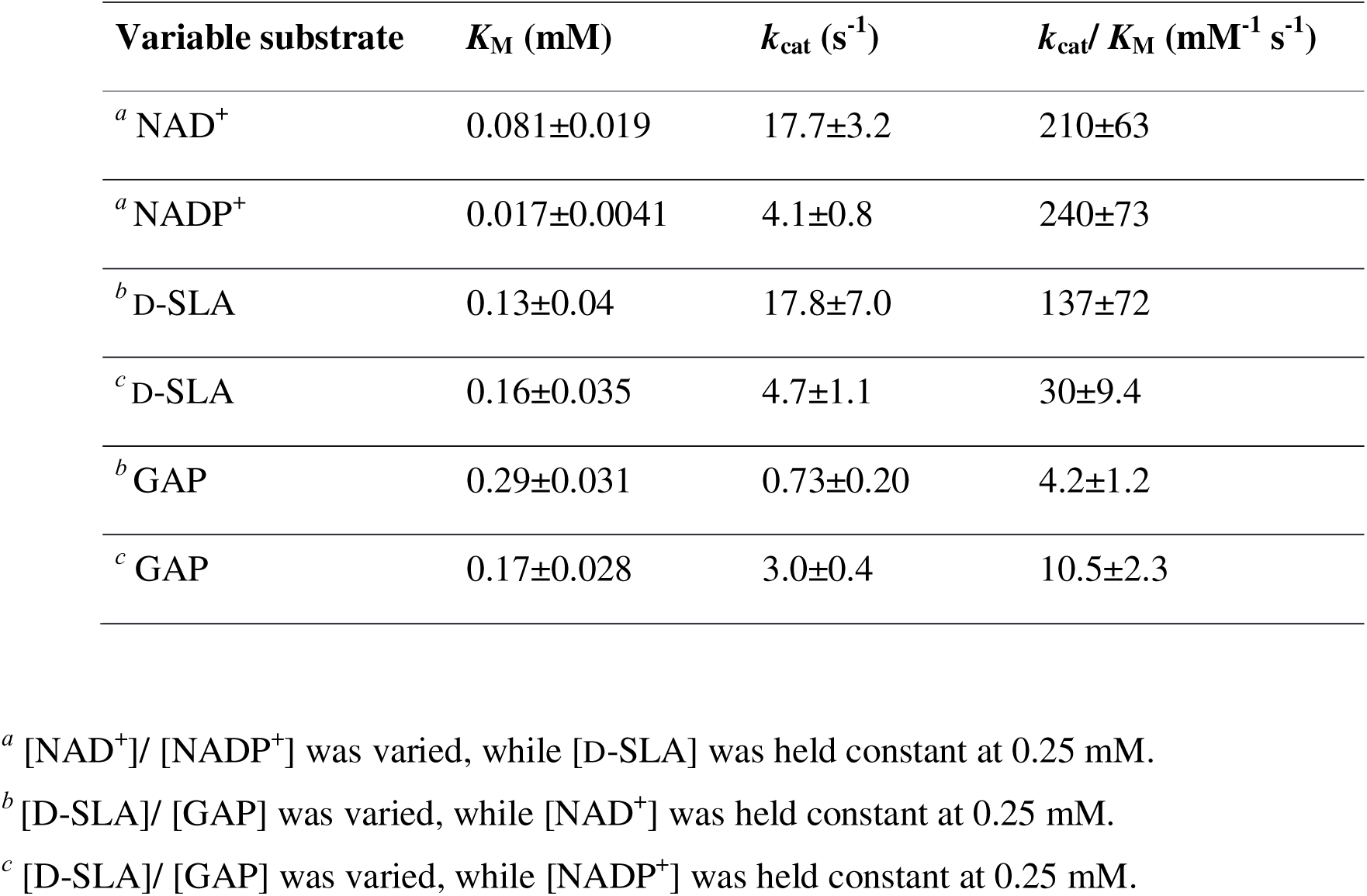
Apparent first order kinetic parameters for *Rl*GabD determined for D-SLA, NAD^+^, NADP^+^ and GAP.

| Variable substrate | $K_M$ (mM) | $k_{cat}$ (s<sup>-1</sup>) | $k_{cat}/K_M$ (mM<sup>-1</sup> s<sup>-1</sup>) |
| --- | --- | --- | --- |
| <sup>a</sup> NAD<sup>+</sup> | 0.081±0.019 | 17.7±3.2 | 210±63 |
| <sup>a</sup> NADP<sup>+</sup> | 0.017±0.0041 | 4.1±0.8 | 240±73 |
| <sup>b</sup> D-SLA | 0.13±0.04 | 17.8±7.0 | 137±72 |
| <sup>c</sup> D-SLA | 0.16±0.035 | 4.7±1.1 | 30±9.4 |
| <sup>b</sup> GAP | 0.29±0.031 | 0.73±0.20 | 4.2±1.2 |
| <sup>c</sup> GAP | 0.17±0.028 | 3.0±0.4 | 10.5±2.3 |

<sup>a</sup> [NAD<sup>+</sup>]/ [NADP<sup>+</sup>] was varied, while [D-SLA] was held constant at 0.25 mM.
<sup>b</sup> [D-SLA]/ [GAP] was varied, while [NAD<sup>+</sup>] was held constant at 0.25 mM.
<sup>c</sup> [D-SLA]/ [GAP] was varied, while [NADP<sup>+</sup>] was held constant at 0.25 mM.

### *Rl*GabD oxidizes GAP and binds reduced NADH analogues

GAP is produced in glycolysis/gluconeogenesis during sulfoglycolytic growth and has a similar structure to SLA. Therefore, we investigated if *Rl*GabD can catalyze the oxidation of GAP. GAP was synthesized from racemic glyceraldehyde-3-phosphate diethyl acetal barium salt.^18, 19^ Apparent Michaelis-Menten kinetics for oxidation of racemic GAP were measured at constant concentration (0.25 mM) of nucleotide (**Fig. 3e,f**). These data reveal that the apparent second order rate constants were similar for NAD^+^ (*k*_cat_/*K*_M_)^app^ = 4.2 s^−1^ mM^−1^ and NADP^+^ (*k*_cat_/*K*_M_)^app^ = 10.5 s^−1^ mM^−1^ with a modest preference for NADP^+^, opposite to that seen for SLA (**Table 1**). As for the kinetics with SLA, inhibition was also observed when the concentration of GAP was higher than 0.25 mM, and so rate data used for Michaelis-Menten analysis was below this limit. The ratio of apparent second order rate constants ((*k*_cat_/*K*_M_)^app^) for SLA and GAP at constant nucleotide concentration reveals that the activity on GAP is approx. 30-fold lower than SLA.

To explore the ability of *Rl*GabD to bind analogues of NADH we synthesized tetrahydro- and hexahydro-NADH by reduction of NADH following the procedure of Dave.^20^ IC_50_ values were measured at constant [SLA] (at *K*_M_ ^SLA^ /10) and constant [NAD^+^] (at *K*_M_ ^NAD+^) (**Fig. S2a,b**). For tetrahydro-NADH, IC_50_ = 28 μM, and for hexahydro-NADH, IC_50_ = 9.1 μM, indicating the latter binds more tightly (**Fig. S2c,d**).

### *Rl*GabD follows a rapid equilibrium ordered kinetic mechanism

*Rl*GabD is a bisubstrate enzyme that acts on two substrates (NAD(P)H and SLA) and produces two products (SL and reduced NAD(P)H) and so its kinetic mechanism is described as Bi-Bi. Such Bi-Bi reactions can occur through non-sequential (Ping Pong) or sequential (ordered, steady-state random, and Theorell-Chance (a special case of ordered reactions where the steady-state level of central complexes is low)) mechanisms.^17^ Analysis of initial rates is a powerful way to distinguish reaction mechanisms. For bisubstrate enzymes with substrates A and B, a plot of 1/υ_0_ versus 1/[A] at various concentrations of substrate B, or 1/υ_0_ versus 1/[B] at various constant concentrations of substrate A can help determine the kinetic mechanism. For a Ping Pong reaction, the plot of 1/υ_0_ versus 1/[A] will afford a series of parallel straight lines with constant slope = *K*_M_(A)/*V*_max_. On the other hand, for a sequential mechanism (ordered or random) the same plot will produce a family of straight lines with slope dependent on the concentration of B and that intersect to the left of the *y* axis, or in the case of a rapid equilibrium ordered mechanism, on the *y* axis itself.

To gain insight into the kinetic mechanism we measured rate data for varying [SLA] at several constant concentrations of NAD^+^, and *vice versa* **(Fig. 4a, d)**. The data were replotted as double reciprocal plots (1/υ_0_ versus 1/[SLA]) **(Fig. 4b, e)**. These primary double reciprocal plots gave a series of intersecting straight lines, consistent with a sequential mechanism. The position of the intersection provides insight into the nature of the sequential or ordered mechanism. The plot of 1/[NAD^+^] versus 1/*V* intersected close to the *y*-axis **(Fig. 4b)**, while the plot of 1/[SLA] versus 1/*V* intersected to the left of the *y*-axis (**Fig. 4e**). While recognizing the difficulty of interpreting whether the intersection of the plot in Fig. 4b is on or close to the *y*-axis, we propose that this data is consistent with a rapid equilibrium ordered mechanism.^17^

**Fig. 4.**
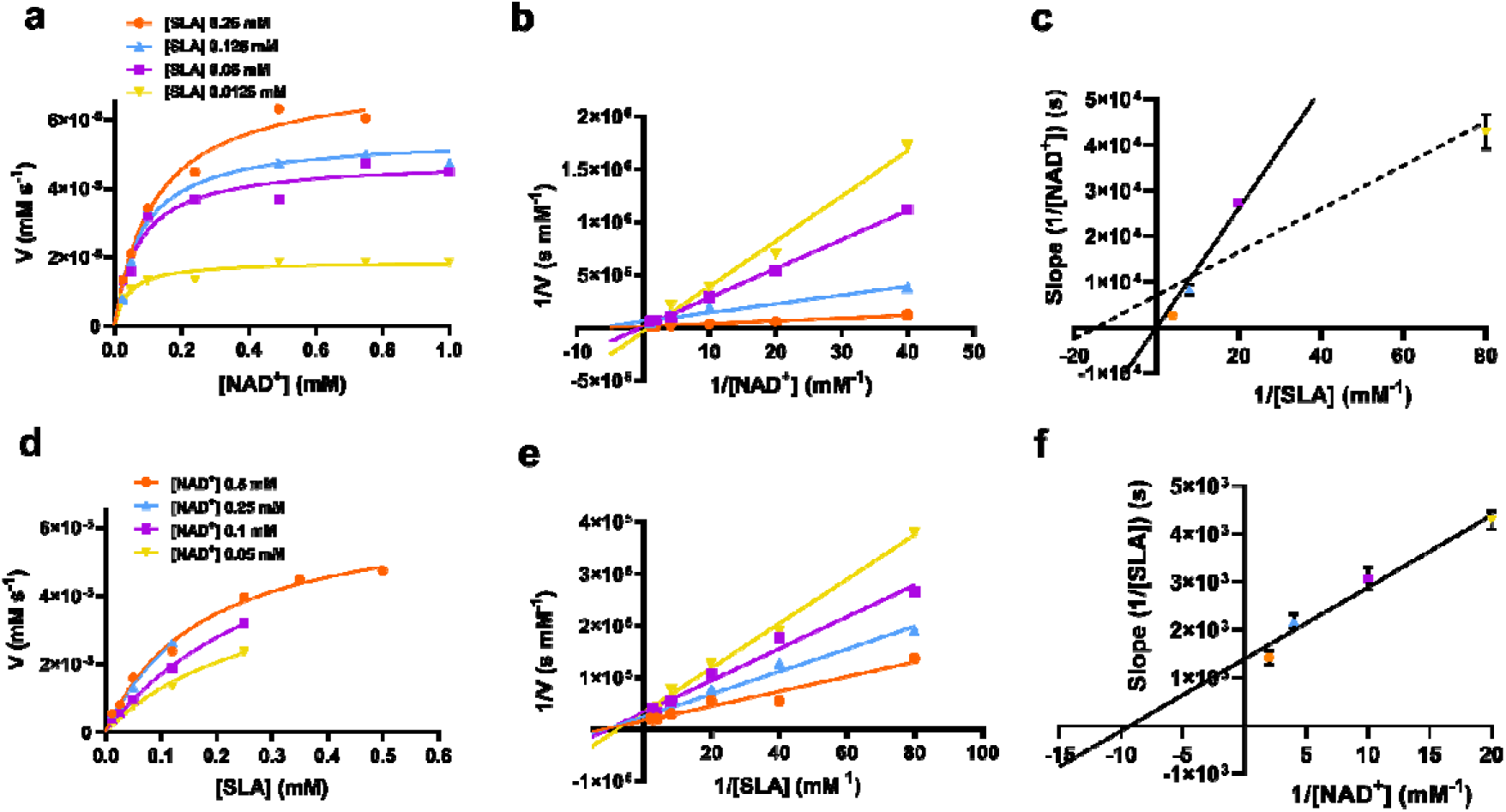
**(a)** Rate data for reactions catalyzed by *Rl*GabD when [NAD^+^] was varied under several different fixed concentrations of D-SLA (0.0125−0.25 mM). **(b)** Double reciprocal plots for the data from (a). **(c)** Secondary plot of slopes from the double reciprocal plot (b). The solid line is fit to the first three data points; the dotted line is fit to all four data points. **(d)** Rate data for reactions catalyzed by *Rl*GabD when [D-SLA] was varied under several different fixed concentrations of NAD^+^ (0.05−0.25 mM). **(e)** Double reciprocal plots for the data from d. **(f)** Secondary plot of slopes from the double reciprocal plot (e). Errors are standard error mean.

Secondary plot analysis involves replotting the slope data from the primary double reciprocal plots. Thus, the slopes of each line in the double reciprocal plot were plotted versus the reciprocal concentrations of the other substrate. For the plot of slopes from the 1/[NAD^+^] versus 1/*V* plot (**Fig. 4c**, see Fig. legend for a more detailed analysis of the possible lines of fit), the line passed through the origin, while for the plot of slopes from the 1/[SLA] versus 1/*V* plot intercepted the *y*-axis above the origin (**Fig. 4f**). Again, recognizing the limits of this graphical approach to determining kinetic mechanism, this data is indicative of a rapid equilibrium ordered reaction, with NAD^+^ binding first to enzyme.^17^

### *Rl*GabD adopts a tetrameric assembly with a classic aldehyde dehydrogenase fold

*Rl*GabD belongs to the family of aldehyde dehydrogenases (ALDHs), which usually exist and function as homodimers and homotetramers.^18^ Size exclusion chromatography with multi-angle laser light scattering (SEC-MALLS) analysis of *Rl*GabD (MW 52,000 Da) showed a solution-state species with molecular weight of approximately 210 kDa (**Fig. S3**). *Rl*GabD thus exists as tetramer in solution, presenting it as a suitable candidate for structural characterisation using cryogenic electron microscopy (cryo-EM).

To define conditions for imaging the complex, we studied the interaction of *Rl*GabD with NAD(H) using nanoscale differential scanning fluorimetry (nano-DSF). nanoDSF uses intrinsic fluorescence to determine the melting temperature of proteins and can aide identification of the formation of protein complexes. nanoDSF revealed a thermal shift (ΔTm) of 4.3 □ for NAD^+^ (and for SLA+NADH, ΔTm of 2.6 □), while NADP^+^ produced a ΔTm of 3.5 □ (and for NADPH ΔTm of 2.5 □) (**Fig. S4**). These results guided experiments to image a binary complex. Thus, cryo-EM grids were optimised and prepared with 1 mg/mL *Rl*GabD pre-incubated with 2 mM NADH. Data collection and refinement statistics of single particle cryo-EM analysis for *Rl*GabD•NADH complex are provided in **Table S1**. A total of 865 micrographs were used for auto-picking. Particles picked from these micrographs were used to generate 2D-class averages, which displayed distinct orientations (**Fig. S5**). Downstream processing and refinement with D2 symmetry gave a final 3D reconstruction of *Rl*GabD at an overall resolution of 2.52 □ at Fourier shell correlation threshold of 0.143 (**Fig. 5a**, **Fig. S6, S7**).

**Fig. 5.**
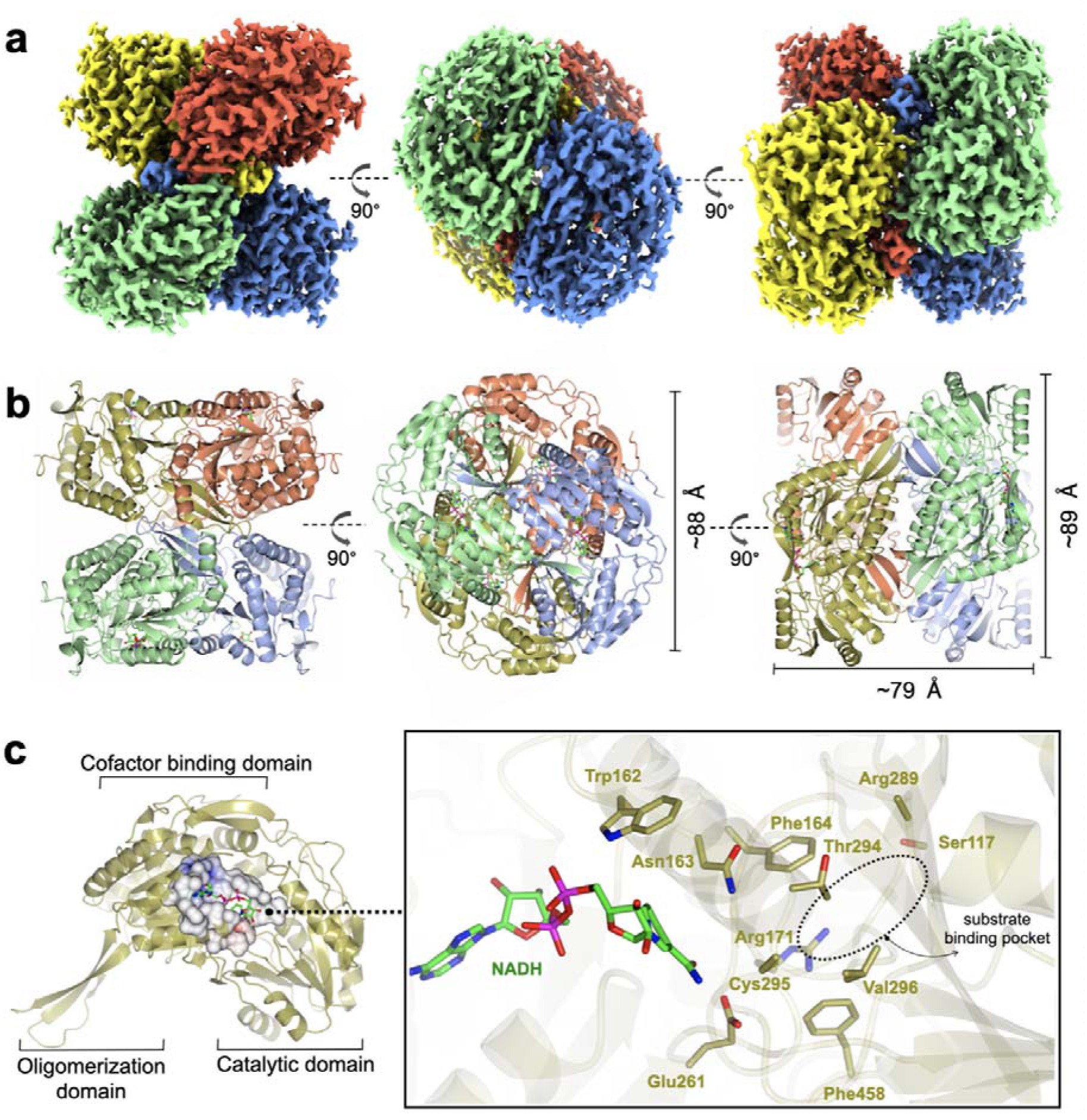
Cryo-EM structure of *Rl*GabD•NADH complex. **(a)** Single-particle cryo-EM reconstruction of the *Rl*GabD•NADH tetramer depicting front and side views. Map is contoured at a threshold of 0.04, and four protomers are coloured in blue, red, green and yellow. **(b)** Quaternary structure of *Rl*GabD tetramer depicting front, top and side views. **(c)** Ribbon representation of an *Rl*GabD monomer showing β-strand oligomerization domain, the NAD-cofactor binding domain, and the catalytic domain. Inset: zoom of active site showing bound NADH molecule and residues lining the active site. The location of the SLA binding pocket is indicated. Cys295 is the predicted catalytic nucleophile, and Glu261 is the predicted general acid/base.

The *Rl*GabD tetramer assembles as a pair of dimers (**Fig. 5a,b**). Each protomer of *Rl*GabD adopts the canonical ALDH class I/II fold with three domains.^19, 20^ Each L-shaped protomer comprises an ⍰/β N-terminal cofactor binding domain [residues 8-133, 153-263], an ⍰/β catalytic domain containing the conserved Cys-Glu dyad [residues 264-476], and a smaller, anti-parallel β-sheet oligomerization domain [residues 134-152, 477-489], which interacts with two other subunits. The *Rl*GabD dimer is formed through domain swapping interactions of the three-stranded oligomerization domain of subunit A with the catalytic domain of partner subunit (B) forming a ten-stranded β-sheet. Oligomerization domains of subunits A+C (and B+D) form extended β-sheets stabilising the final pair-of-dimers assembly.

### A binary *Rl*GabD•NADH complex reveals the active site architecture

The *Rl*GabD**•**NADH complex shows the binding mode and positioning of the cofactor. Poor density was evident for the second phosphate and of the ribosyl-nicotinamide group (**Fig. S9**). However, this still allowed NADH to be modelled with confidence, and showed that NADH is bound in an extended mode, as seen in *E. coli* SSADH GabD,^21^ with the nicotinamide group protruding into the active site located in the cleft between the two major domains (**Fig. 5c**). The nicotinamide ring sits in close vicinity to the catalytic dyad, and the adenine ring occupies a hydrophobic pocket lined by Ala 219, Leu224, Val243, Trp246, and Leu247. Based on the distance between heteroatoms, the 2’-OH of the adenosine of NADH forms hydrogen bonds with Lys186 (2.5 □) and a water molecule, which in turn engages in hydrogen bonding interactions with Ser189 (2.6 □). The 3’-OH of adenosine is hydrogen-bonded to Lys186 (3 □) and the backbone carbonyl of Thr160 (2.4 □). The pyrophosphate group of NADH forms hydrogen-bonding interactions with the backbone carbonyl and hydroxyl group of Ser240. Some additional density is seen at the active site Cys295 residue, possibly indicating oxidation and weaker side-chain density of Glu261 owing to radiation damage or cryo-EM density weakness of negatively charges of carboxylate groups.^22, 23^

The poor density in the core of the bound NADH may reflect multiple binding modes of the cofactor. At least two discrete conformations have been reported for the nicotinamide ring in members of ALDH class I/II families. *Rl*GabD shares high sequence and structural and functional similarities with representative ALDH members such as *E. coli* SSADH (PDB: 3JZ4, core RMSD of 0.65 □ and 61% sequence ID),^21^ *E. coli* lactaldehyde DH (PDB: 2ILU, RMSD 1.3 □ and 35% sequence ID),^24^ and the reduced form of human SSADH (PDB: 2W8R, RMSD 0.76 □ and 55% sequence ID).^25^ Structural comparison of the *Rl*GabD•NADH complex with *E. coli* SSADH and lactaldehyde DH demonstrates the two discrete ‘in’ and ‘out’ cofactor conformations (**Fig. S10**). The *Rl*GabD•NADH complex displays the catalytically-relevant ‘in’ conformation with the nicotinamide ring pointing into the active site, and C4 of nicotinamide approx. 6.7 □ from catalytic Cys295. Further, the 2’-phosphate binding residues Ser179 and Lys182 (*E. coli* SSADH numbering) are conserved in *Rl*GabD, contributing the dual cofactor specificity [NAD(P)H] of *Rl*GabD.

To propose active site residues involved in catalysis we compared the sequence alignment of *Rl*GabD with the NADP-dependent non-phosphorylating glyceraldehyde 3-phosphate dehydrogenase (GAPN) from *Streptococcus mutans* (**Fig. S15**).^26, 27^ Like *Rl*GabD, GAPN operates through an ordered sequential mechanism in which the cofactor binds first.^26^ The catalytic mechanism for oxidation of GAP by GAPN has been described in detail and involves two main steps.^27^ In the first step, nucleophilic addition of Cys302 to the aldehyde of GAP forms a hemithioacetal oxyanion, which is stabilized by an ‘oxyanion hole’ formed from the terminal NH_2_ of Asn169 and the backbone N-H of Cys302. The hemithioacetal oxyanion is activated to transfer hydride to NADP, forming an acyl enzyme and NADPH. In the second step, Glu268 acts as general base to assist the nucleophilic addition of water to the acyl enzyme, forming a tetrahedral intermediate oxyanion, which eliminates Cys302 to give the product, 3-phosphoglycerate. All of the residues involved in GAPN catalysis are conserved with *Rl*GabD andthe 3D structure reveals that they are in an appropriate position adjacent to the nicotinamide headgroup of NADH to adopt similar roles (**Fig. 5**). Overlay of the 3D structures of *Rl*GabD and that of the covalent thioacyl adduct of the Glu268Ala mutant of GAPN^27^ reveals spatial conservation of the bases (*Rl*GabD Glu261, GAPN Glu268Ala) and nucleophiles (*Rl*GabD Cys295, GAPN Cys284) (**Fig. 6**). Thus, we propose that Cys295 is the catalytic nucleophile, Glu261 is the general base, and the oxyanion hole is formed from Asn163 and the backbone NH of Cys295. We probed the importance of Cys295 and Glu261 for catalysis by site-directed mutagenesis. The relative activity versus wildtype for Cys295Ala was 1/120,000 and for Glu261Ala was 1/63,000 at 0.5 mM NAD^+^ and 0.25 mM D-SLA (activity was undetectable with 0.5 mM NADP^+^ and 0.25 mM D-SLA). These values approach the limits of site directed mutagenesis because of the complications of translational misincorporation by the heterologous host *E. coli*. Thus, both the Cys295Ala and Glu261Ala variants are severely disabled catalysts, consistent with their critical roles in catalysis.

**Fig. 6.**
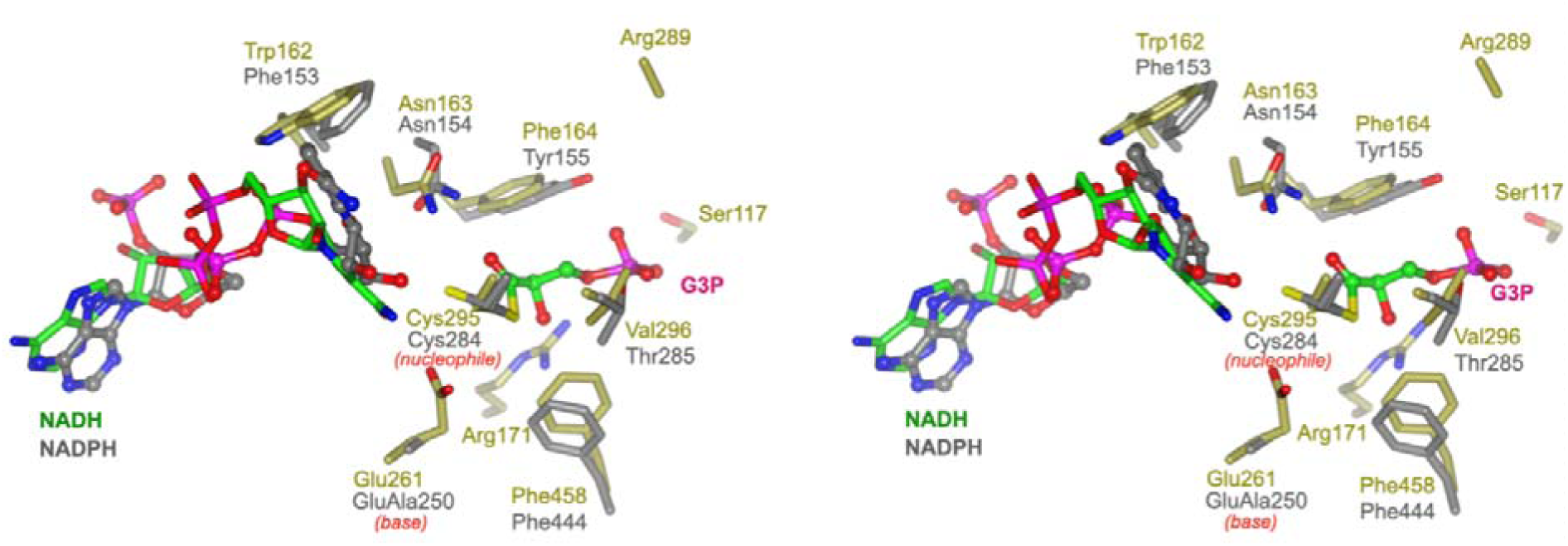
Conservation of active site residues of *Rl*GabD with glyceraldehyde 3-phosphate dehydrogenase. Stereoview of *Rl*GabD•NADH (gold) and NADPH complex of the acyl enzyme intermediate formed on the Glu268Ala mutant of glyceraldehyde 3-phosphate dehydrogenase from *Streptococcus mutans* (PDB code: 2ESD, grey). The structures align with an RMSD of 1.26 over 453 residues. Cys295 is the predicted catalytic nucleophile, and Glu261 is the predicted general acid/base.

### Prediction of the SLA binding pocket in SLADH enzymes

To predict the location of the SLA binding pocket, we computed the interior cavities and channels in the *Rl*GabD•NADH tetramer using the CASTp server 3.0^28^ (**Fig. S11, S12**). This predicted a hydrophilic, active-site pocket with solvent-accessible area of approx. 283 [^2^ and volume of 184 [^3^ (reported as Richard’s solvent accessible surface area/volume^29^), with helix 173 [residues 106-121] and surface loop [residues 449-458] forming the mouth of the opening. The base of this cavity is buried in the cleft between the two domains and adjoins the conserved nucleophile Cys295. Superposition of the predicted SLA binding pocket computed by CASTp with the 3D structure of human SSADH (in which the catalytic nucleophile Cys340 was converted to Ala) with bound succinate semialdehyde (SSA) (PDB 2W8Q)^25^ shows SSA occupies this cavity in an orientation where aldehyde group directly points towards the nucleophile Cys295 of *Rl*GabD, suggesting a similar orientation for sulfolactaldehyde (SLA). The carboxylate group of SSA is H-bonded to Arg213, Arg334 and Ser498 (hSSADH numbering). These residues are incompletely conserved with SLADH enzymes. *Rl*GabD and several other SLADH candidates including *P. putida*, *Arthrobacter* sp., and *Desulfovibrio* sp. contain Arg residues at equivalent positions (*Rl*GabD: Arg171 and Arg289), but *B. megaterium* and *B. urumqiensis* have Arg213 is replaced by His, and Arg334 by Asn. The multiple sequence alignment at hSSADH position Ser498 is poorly aligned with insertions/deletions, and no clear consensus. We propose Arg171/Arg289 as sulfonate binding residues in *Rl*GabD and some SLADH enzymes, with the equivalent positions as His/Asn in other SLADH enzymes fulfilling a similar role. To explore whether other residues are associated with the Arg171/Arg289 pair in *Rl*GabD, and the His164/Asn280 pair in *B. megaterium* SlaB we conducted coevolution analysis using CoeViz.^30^ Using the multiple sequence alignment of 158 putative SLADH enzymes (*vide infra*) we identified a clique of 16 residues (including the above pairs) that independently co-evolve with the above pairs (**Fig. S13**). Mapping of these coevolving cliques onto the cryo-EM structure of *Rl*GabD and the AlphaFold2^31, 32^ model of *Bacillus megaterium* SlaB identified a tripeptide sequence of partially conserved residues in proximity to the proposed SLA binding pocket that contain the oxyanion hole stabilizing residue, namely Trp162-Asn163-Phe164 in *Rl*GabD and Phe155-Asn156-Val157 in *B. megaterium* SlaB.

Human SSADH (hSSADH) shares a similar fold and catalytic residues with *E. coli* GabD SSADH and *Rl*GabD SLADH, including the catalytic cysteine (Cys340 in hSSADH). hSSADH contains a second cysteine (C342) two residues downstream in a redox active mobile loop that can engage in a disulfide bond with the nucleophilic cysteine. Oxidation to the disulfide results in hSSADH adopting an ‘closed’ conformation, while reduction to cysteine causes loop movement and a ‘open’ conformation (**Fig. S14**). The second cysteine residue is not conserved in *E. coli* SSADH nor some SLADH enzymes (e.g. *Arthrobacter* spp., *B. urumqiensis*, and *B. megaterium*), but is present in *Rl*GabD and SLADHs from *Desulfovibrio* sp., and *P. putida* (**Fig. S15**). In the *Rl*GabD•NADH structure, the catalytic loop of *Rl*GabD adopts the ‘open’ conformation, with the two cysteine residues 8.4 Å apart and the catalytic dyad Cys295/Glu261) poised for catalysis (**Fig. S14**). It is unknown whether Cys295/297 in bacterial SLADH proteins undergo comparable oxidation and associated loop movement as seen for hSSADH.

### Sequence similarity network analysis reveals the taxonomic range and functional distribution of SLA dehydrogenases in the pathways of sulfoglycolysis

SLA dehydrogenases occur in several sulfoglycolytic pathways: sulfo-ED, sulfo-SFT, and sulfo-EMP pathways, as well as within DHPS degradation pathways. To explore the distribution and evolution of SLA reductases we performed sequence similarity network (SSN) analysis^33^ using the EFI enzyme similarity tool (EFI-EST).^34, 35^ Using the individual SLA dehydrogenase sequences from seven experimentally-verified sulfoglycolytic organisms (*B. urumquiensis*, *P. putida*, *R. leguminosarum*, *H. seropedicae*, *B. megaterium*, *Arthrobacter* sp. AK01, *B. aryabhattai*) and one DHPS degrading organism (*Desulfovibrio* sp. DF1) we separately conducted BLASTp searches and combined the results to obtain a total of 158 sequences with >37-64% sequence identity to the search queries. To visualize and study the distribution of these sequences, we used SSNs. Initially, we explored the construction of SSNs using different alignment scores (**Fig. S16**). At alignment score 75, the sequences form a single cluster; at alignment score 100 two clusters; while in the range 125-150, the SSN breaks into three clusters that almost perfectly separate the three main Phyla: Actinobacteria, Firmicutes and Proteobacteria, with the three Chlorflexi members, the sole Candidatus Dormibacteraeota member and one spurious Firmicutes member clustering with Actinobacteria. At even higher alignment threshold (175), the Chloroflexi members segregate, but the Actinobacteria fragment and the SSN spawns many singletons that limits its utility. The SSN generated at alignment score 150 (corresponding to minimum identity >53%; **Fig. 7a**) was coloured based on the pathway encoded by the proposed function of the gene cluster in which the SLA dehydrogenase gene was located (**Fig. 7b**). The sulfo-EMP pathway organisms are limited to Actinobacteria; sulfo-EMP2 pathway organisms are mainly limited to Firmicutes but with several members within the Actinobacteria; while hybrid sulfo-EMP pathways comprised of various combinations of genes from the sulfo-EMP and sulfo-EMP2 pathways are more broadly distributed across Actinobacteria, Firmicutes, and Proteobacteria. Sulfo-ED organisms occur mainly within proteobacteria but with several members within the Actinobacteria. Sulfo-SFT organisms are mainly Firmicutes but with membership of two Chloroflexi and one Candidatus Dormibacteraeota. Finally, DHPS degradation pathways are mainly limited to Actinobacteria with a sole Proteobacteria representative. We highlight the Proteobacteria member *Ensifer* sp. HO-A22 contains both sulfo-ED and DHPS degrading gene clusters, suggesting that this organism achieves the complete biomineralization of SQ to sulfite. A tree showing the phylogenetic relationships based on 16S ribosomal RNA sequences between bacteria that contain SLADH genes within sulfoglycolytic gene clusters is shown in **Fig. S17**.

**Fig. 7.**
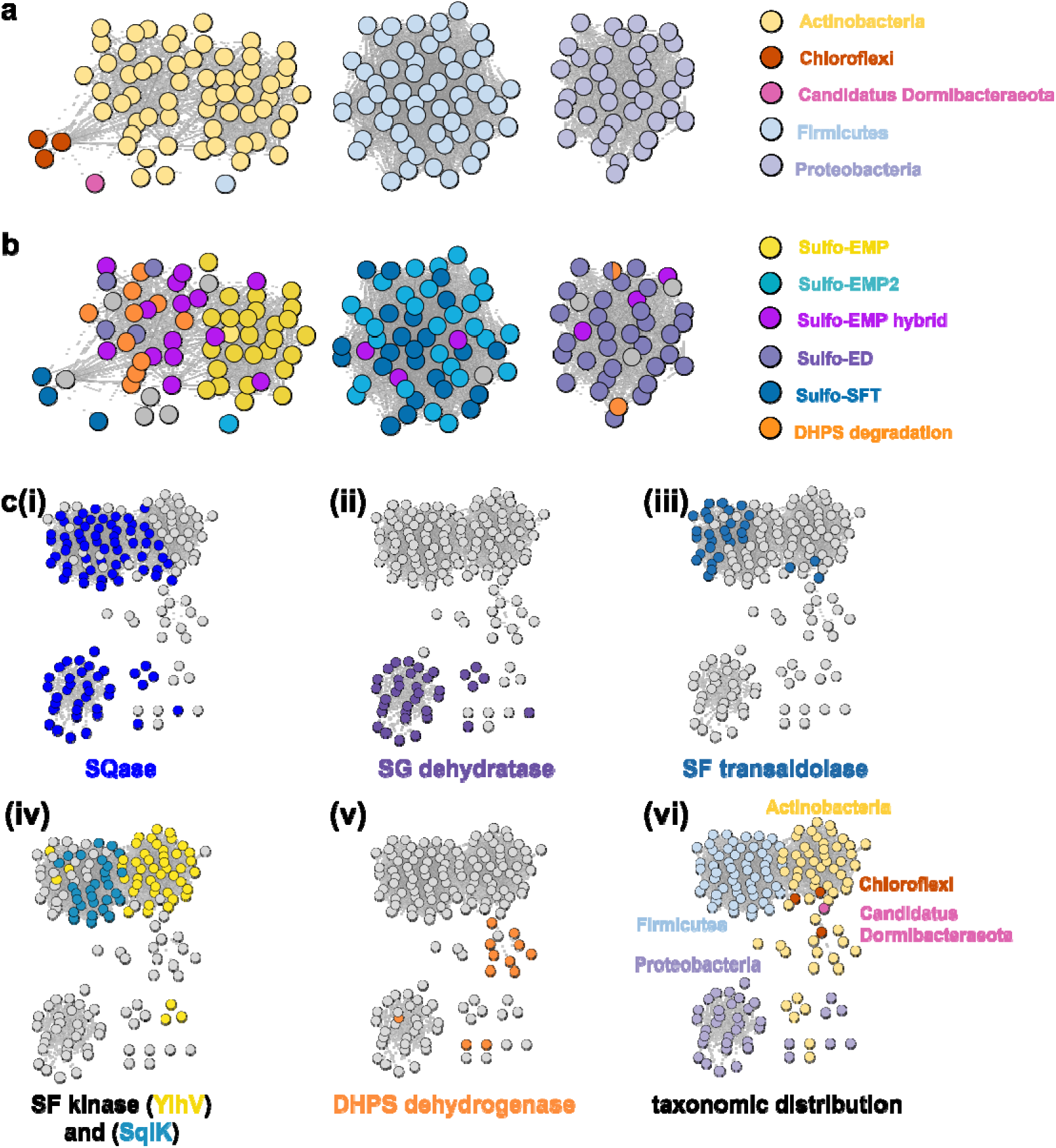
SSN of SLA dehydrogenase proteins (family PF00171) showing distribution in Actinobacteria, Firmicutes, Proteobacteria, Chloroflexi and Candidatus Dormibacteraeota. Nodes are individual SLADH proteins that are coloured according to: **(a)** occurrence within indicated SQ or DHPS degradation pathways, or (**b)** distribution across five phyla. **(c)** Genome neighborhood similarity network (GNSN) of SLA dehydrogenase proteins. Each node corresponds to an SLA dehydrogenase ortholog in the family PF00171. Nodes are colored according to the presence of genes encoding SQ or DHPS degradation enzymes within a ±10-ORF window of the gene encoding SLA dehydrogenase. Each SQ or DHPS degradation enzyme corresponds to a specific cluster in the SSNN in **Fig. S18**. Edges connect nodes that share >4 isofunctional genes in their genome neighborhood. Nodes are coloured in each panel if a specific enzyme belonging to a PFAM is found in the genome neighborhood of the SLA dehydrogenase otholog: **(i)** SQase (PF01055); **(ii)** SG dehydratase (PF00920); **(iii)** SF transaldolase (PF00923); **(iv)** YihV-type SF kinase (PF00294), SqiK-type SF kinase (PF00365); **(v)** DHPS dehydrogenase (PF03446); **(vi)** nodes are coloured according to the Phyla of the host organism.

We used the sequences from the SSN to identify the genes that flank the 158 SLADH genes in the genomes of the host organisms. Using the EFI-EST tools, we identified 1287 gene neighbours located ± 10 ORF from the query SLADH sequences. These were analysed by creation of a sequence similarity network of neighbors (SSNN) into isofunctional proteins that were assigned a function based on manual inspection (**Fig. S18**). To organize and visualize the sulfoquinovose and DHPS degrading gene clusters we constructed a genome neighborhood similarity network (GNSN) using the EFI-GNT tool (**Fig. 7c**). In this network each node corresponds to a single SLA dehydrogenase protein that is connected by an edge to another SLA dehydrogenase if they share >4 isofunctional genes in their genome neighborhood. The GNSN shows that SQase proteins are encoded in the gene clusters for most sulfoglycolytic organisms, with the exception of some sulfo-EMP organisms, consistent with the role of SQases as a gateway to sulfoglycolysis through cleavage of SQ-glycosides (**Fig. 7c(i)**).^36, 37^ Characteristic enzymes encoded by sulfo-EMP (SF kinase YihV), sulfo-EMP2 (SF kinase SqiK), sulfo-ED (SG dehydratase), sulfo-SFT (SF transaldolase) and DHPS degradation (DHPS dehydrogenase) pathways distribute across the GNSN into clusters (**Fig. 7c(ii-v)**). The sulfoglycolytic clusters are mutually exclusive to the DHPS degrading clusters, except for *Ensifer* sp. HO-A22, which occurs within the main sulfo-ED cluster. When the GNSN was coloured for the five phyla identified in the SSN (Actinobacteria, Firmicutes, Proteobacteria, Chloroflexi, and Candidatus Dormibacteraeota) (**Fig. 6c(vi)**), we observed coloured clusters that recapitulated the taxonomic clustering of the SSN of SLADH sequences in **Fig. 6a**.

## Conclusions

SLADH enzymes catalyze the oxidation of SLA to SL, the final step of sulfoglycolytic pathways that lead to excretion of SL,^4^ and the second step in the oxidation of DHPS to SL in *Desulfovibrio* sp.,^5^ which activates this substrate for sulfur-carbon bond scission in the DHPS degradation pathway to produce sulfite and pyruvate. Our data demonstrates that *Rl*GabD has dual cofactor activity, and a 30-fold preference for oxidation of SLA versus the structurally-related phosphate analogue glyceraldehyde phosphate, a key intermediate in glycolysis/gluconeogenesis. The weak activity on GAP will lead to production of 3-phospho-D-glycerate, an intermediate in glycolysis, and thus this low-level activity likely has little consequence for cellular metabolism. Nonetheless, the activity of SLA dehydrogenase on GAP stands in contrast to *E. coli* SLA reductase, which had no detectable activity on GAP.^7^

Similar to the well-characterized GAP dehydrogenase from *S. mutans*,^26^ our data suggests that *Rl*GabD uses a rapid equilibrium ordered mechanism, in which NAD(P)^+^ is the first substrate to bind. Knowledge of reaction order, a large change in protein melting temperature upon binding NADH, and the identification of a tetramer in the solution state, guided our approach to determining the 3D structure of the *Rl*GabD•NADH complex using cryo-EM. This complex revealed sequence and spatial conservation of amino acid residues involved in catalysis, and allows proposal of a mechanism for catalysis (**Fig. 8**). Binding of NAD(P)^+^, and then SLA gives the Michaelis complex. In the first step, nucleophilic addition of Cys295 to the aldehyde of SLA forms a hemithioacetal oxyanion, stabilized by an ‘oxyanion hole’ formed from the terminal NH_2_ of Asn163 and the backbone N-H of Cys295. The hemithioacetal oxyanion is activated to transfer hydride to NAD(P)^+^, forming an acyl enzyme and NAD(P)H. In the second step, Glu261 provides general base catalysis, assisting the nucleophilic addition of water to the acyl enzyme, forming a tetrahedral intermediate oxyanion, which eliminates Cys295 to give SL. Based on their proximity to the active site, we propose that Arg171-Arg289 comprise the sulfonate binding residues in *Rl*GabD, and the first and last residues within the tripeptide sequence Trp162-Asn163-Phe164 (containing the oxyanion stabilizing residue) comprise additional SLA binding residues. Arginine residues are common in a wide range of other sulfonate binding proteins and enzymes from various sulfoglycolytic pathways.

**Fig. 8.**
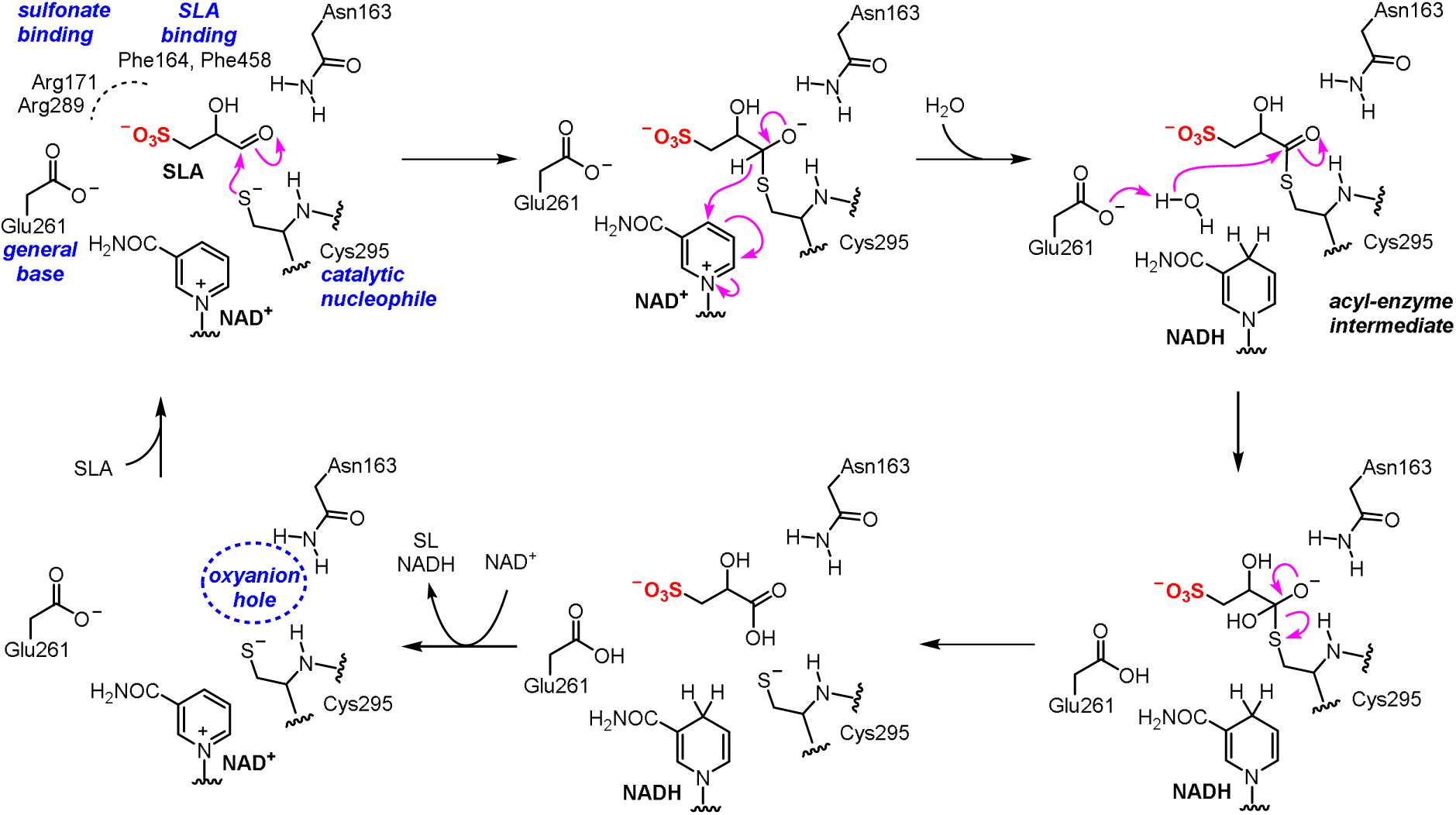
Proposed mechanism and active site residue roles for *Rl*GabD SLA dehydrogenase.

Because SL as an endproduct of sulfoglycolysis, a nutrient for SL degrading bacteria, and an intermediate in DHPS degradation, the oxidation of SLA to SL catalyzed by SLADH is an important step in the breakdown of the C6-organosulfonate sulfoquinovose and the C3-organosulfonate DHPS. Sulfoglycolytic gene clusters containing genes encoding SLADH enzymes are distributed across Actinobacteria, Firmicutes, Proteobacteria, Chloroflexi, and Candidatus Dormibacteraeota, while DHPS degradation gene clusters containing SLADH homologues are limited to Proteobacteria and Actinobacteria. The present work provides a structural and biochemical view of SLADH enzymes that complements our knowledge of SLA reductases, and enriches our understanding of a critical step in the organosulfur cycle.

## Data availability

The supporting information includes Experimental and additional details on enzyme kinetics (Fig. S1-2), protein biochemistry (Fig. S3-S4), Cryo-EM (Fig. S5-S9), 3D structural data (Fig. S10-S12), bioinformatics (Fig. S13-S18), and structural statistics (Table S1)

## Author contributions

G.J.D. and S.J.W. conceived the project. N.M.S. and M.S. performed molecular biology and protein purification. M.S., Z.A., and R.M. prepared the cryo-EM grids and performed the image acquisition and processing. J.L. and A.A. synthesized SLA and inhibitors, and conducted enzyme kinetics. J.N.B. helped with the image acquisition, processing, and model building. J.L. and M.S. conducted bioinformatics. J.L., M.S., A.A., E.D.G.-B., J.N.B., G.J.D. and S.J.W. analysed the data and wrote the manuscript.

## Acknowledgements

This work was supported by the Australian Research Council (DP210100233, DP210100235), the Biotechnology and Biological Sciences Research Council (BB/W003805/1), the UKRI Future Leader Fellowship Program (MR/T040742/1), and the Royal Society for the Ken Murray Research Professorship to G.J.D. and the associated PDRA funding (RP\EA\180016) for RWM. J.L. is supported by the China Scholarship Council. E.D.G.-B. acknowledges support from The Walter and Eliza Hall Institute of Medical Research, National Health and Medical Research Council of Australia (NHMRC) project grant GNT2000517, the Australian Cancer Research Fund, and the Brian M. Davis Charitable Foundation Centenary Fellowship. We thank the Wellcome Trust for funding the Glacios electron microscope (grant number 206161/Z/17/Z) and Dr Johan Turkenburg and Sam Hart for assistance with cryo-EM data collection. We also acknowledge Dr. Andrew Leech at the University of York Bioscience Technology Facility for assistance with SEC-MALLS analysis, and Arashdeep Kaur for assistance with bioinformatics.

## Abbreviations

DHPS: 2,3-dihydroxypropanesulfonate
ED: Entner-Doudoroff
EMP: Embden-Meyerhof-Parnas
GAP: glyceraldehyde-3-phosphate
SL: sulfolactate
SLA: sulfolactaldehyde
SQ: sulfoquinovose
SSA: succinate semialdehyde
NAD(P)H: reduced nicotinamide adenine dinucleotide (phosphate)

