## Supplementary material file for "Molecular basis of sulfolactate synthesis by sulfolactaldehyde dehydrogenase from *Rhizobium leguminosarum*"

**Experimental section**

**Heterologous expression and protein purification**

The sequence encoding RlGabD (WP_017967313.1) was amplified from *Rhizobium leguminosarum* SRDI-565 genomic DNA by PCR using the primers 5’-TTACTCATATGCCATCCAACTATGACAGC-3’ and 5’-TCAGACTCGAGGTTCTGCGTCAACCCGGCATC-3’. The amplicon was cloned into the MCS of pET-29b(+) (Novagen) using the *NdeI* and *XhoI* sites, and sequence-verified by Sanger sequencing. This plasmid was transformed into chemically competent *E. coli* BL21(DE3) cells and starter cultures grown in LB-medium (5 mL) containing 100 µg mL^-1^ kanamycin for 18 h at 37 °C with shaking at 200 r.p.m. 1 L volume cultures were inoculated with the starter culture (5 mL) and incubated at 37 °C, with shaking at 200 r.p.m. until an A_600_ of 0.6-0.8 was reached. *Rl*GabD expression was induced by addition of IPTG (0.5 mM) and shaking continued overnight at 18 °C at 200 r.p.m. The cells were harvested by centrifugation at 5000 *g* for 20 min and the pellet resuspended in 50 mM sodium phosphate buffer pH 7.4, containing 500 mM NaCl and 30 mM imidazole. Cells were disrupted by ultrasonication for 3 × 5 min, 30 s on; 30 s off cycles, and the suspension was centrifuged at 50,000 *g* for 30 min to yield a clear lysate. The C-terminal His_6_-tagged protein was purified by immobilized-metal affinity chromatography (IMAC) using a Ni-NTA column, followed by size exclusion chromatography (SEC). For IMAC, the lysate was loaded onto a pre-equilibrated Ni-NTA column, followed by washing with a load buffer (50 mM Tris, 500 mM NaCl, 30 mM imidazole pH 7.5). The bound protein was eluted using a linear gradient with buffer containing 300 mM imidazole. *Rl*GabD fractions were pooled, concentrated and loaded onto a HiLoad 16/600 Superdex 75 gel filtration column pre-equilibrated with 50 mM Tris, 300 mM NaCl pH 7.5 buffer. The protein was concentrated using a Vivaspin® 6 with a 30 kDa MW cut-off membrane, to a final concentration of 8 mg mL^-1^ for structural studies.

**Generation of active-site variant constructs**

The active-site variant constructs, *Rl*GabD Glu261Ala and *Rl*GabD Cys295Ala, were generated with the Q5 site-directed mutagenesis kit (New England Biolabs) using the primers 5’-cgTTGGGCGGCAATGCGC-3’ and 5’-cCAGCGAGAGGCGCTTG-3’ for Glu261Ala and 5’-CGGCCAGACCgcgGTTTGCGCCA-3’ and 5’-GCATTGCGGAATTTGGAG-3’ for Cys295Ala, respectively (lower-case letters indicate the mutated sequence). Mutagenesis was verified by DNA sequencing and the proteins were purified by the same method used for the wild-type protein.

**SEC-MALLS analysis**

Experiments were conducted on a system comprising a Wyatt HELEOS-II multi-angle light scattering detector and a Wyatt rEX refractive index detector linked to a Shimadzu HPLC system (SPD-20A UV detector, LC20-AD isocratic pump system, DGU-20A3 degasser and SIL-20A autosampler). Work was conducted at room temperature (20±2 °C). Solvent was 0.2 µm filtered before use and a further 0.1 µm filter was present in the flow path. The column was equilibrated with at least 2 column volumes of buffer (50 mM NaPi, 300 mM NaCl pH 7.4) before use and flow was continued at the working flow rate until baselines for UV, light scattering and refractive index detectors were stable. Sample injection volume was 100 µL *Rl*GabD at 6 mg mL^-1^ in 50 mM Tris buffer, 300 mM NaCl pH 7.4 containing 2 mM NADH; Shimadzu LC Solutions software was used to control the HPLC and Astra V software for the HELEOS-II and rEX detectors. The Astra data collection was 1 min shorter than the LC solutions run to maintain synchronisation. Blank buffer injections were used as appropriate to check for carry-over between sample runs. Data were analysed using the Astra V software. MWs were estimated using the Zimm fit method with degree 1. A value of 0.182 was used for protein refractive index increment (dn/dc).

**Nanoscale Differential Scanning Fluorimetry (nanoDSF)**

NanoDSF studies were performed on a Prometheus NT.48 (NanoTemper). Data recording and initial analysis was performed with PR.ThermControl software. All protein samples were at 2 mg.ml-1 in 50 mM Tris, 150 mM NaCl at pH 7.4, with a 15 μl capillary load per sample. Temperature was ramped from 15 °C to 95 °C, at 1.0 °C/min with 10% excitation power. Experiments were performed in duplicate.

**Synthesis of SLA and determination of concentration**

SLA was synthesized as a solution in water as reported, at a nominal concentration of 109 mM.^17^ The concentration of the SLA solution was measured by quantitative ^1^H NMR spectroscopy using a calibration curve of methyl β-glucoside to determine the instrument sensitivity. The calibration curve was generated by measuring the absolute integration of H1 (δ 4.25 ppm in D_2_O) of methyl β-glucoside at 100, 50, and 10 mM. The absolute integration of H1 was 15738, 7753.06, 1249.15, respectively **(Fig. S1A)**. The SLA stock solution was diluted 1:1 with D_2_O. The absolute integration of H2 (δ 3.79 ppm in D_2_O) of SLA in the ^1^H NMR spectrum was 8647.6. The calculated concentration of SLA was 110 mM, which was used for all further calculations.

**Measurement of consumption of SLA by *Rl*GabD**

The production of NADH from oxidation of SLA catalyzed by *Rl*GabD was monitored using UV-Vis spectrophotometer at 340 nm, in triplicate. The reaction was carried out in 30 mM Tris buffer, 30 mM KCl pH 7 at 30 °C [NAD^+^] = 1.5 mM, [DL-SLA] = 1 mM, and [*Rl*GabD] = 15.8 nM. Phosphate buffer was not used as it has been shown to have a negative impact on NADH stability.^24^ The reaction appeared complete after 40 min **(Fig. S1B)**. More *Rl*GabD was added to a final [*Rl*GabD] = 205 nM, and the reaction was monitored for 100 min with no further increase in absorbance. The extinction coefficient used for NADH was 6363 M^–1^ cm^–1^. Error is standard error mean.

**Michaelis-Menten kinetic analysis of *Rl*GabD**

Michaelis-Menten kinetic analysis were performed for SLA, GAP, NAD^+^ and NADP^+^ under pseudo first-order conditions in which the concentration of one substrate was varied while the other was held constant, and *vice versa*. The production of NADH/NADPH from oxidation of SLA catalyzed by *Rl*GabD SLA dehydrogenase was monitored using a UV-Vis spectrophotometer at 340 nm. For the Michaelis-Menten kinetics of NAD^+^/NADP^+^, reactions were conducted in 0.1% BSA 30 mM Tris buffer pH 7.12 at 30 °C with 3 nM *Rl*GabD, constant 0.25 mM D-SLA and varying concentrations of NAD^+^/NADP^+^ (0.05-1 mM). For the Michaelis-Menten kinetics of D-SLA, constant 0.25 mM NAD^+^/NADP^+^ and varying concentrations of D-SLA (0.025-0.25 mM) were used. For the Michaelis-Menten kinetics of racemic GAP, constant 0.25 mM NAD^+^/NADP^+^ and varying concentrations of GAP (0.05-0.25 mM) were used. The apparent kinetic parameters, *k*_cat_, *K*_M_, and *k*_cat_*/K*_M_ were calculated using the Prism 9 software package (GraphPad Scientific Software) **(Table 1)**. No reaction was observed for >0.5 mM [D-SLA] and >0.25 mM [NADP^+^]. This appears to be a result of inhibition.

The activity of Cys295Ala and Glu261Ala variants were measured at 0.5 mM NAD^+^ and 0.25 mM D-SLA, or 0.5 mM NADP^+^ and 0.25 mM D-SLA) in 0.1% BSA 30 mM Tris buffer pH 7.12 at 30 °C. The concentration of each variant enzyme was 3000 nM.

**Cryo-EM 3D-structure of *Rl*GabD•NADH**

A frozen aliquot of *Rl*GabD was prepared at 4 mg/mL and complexed with 2 mM NADH for screening. R1.2/1.3 300 mesh UltrAuFoil gold grids (Quantifoil) were glow-discharged for 3 min at 20 mAmp/0.38 mBar. A total of 2.5 µL of this sample was applied to glow-discharged UltrAuFoil gold grids, which were subsequently blotted for 2 s with a blot force of 10, then were plunged into liquid ethane cooled by liquid nitrogen. Plunge-freezing was performed using a Vitrobot Mark IV (Thermo Fisher Scientific) at 100% humidity and 22 °C. EER formatted movies of *Rl*GabD•NADH complex were acquired on the Glacios microscope (Thermo Fisher Scientific), housed in the York Structural Biology Laboratory. The microscope was operated at an accelerating voltage of 200 kV with a Falcon 4 direct electron detector. The *Rl*GabD•NADH dataset was acquired at a dose rate of 2.98 electrons per pixel per second, and a pixel size of 0.574 Å; target defocus values were -2 to -0.7 μm. The autofocus function was run every 10 μm, and the dataset was collected with a total dose of 50 electrons per Å^2^.

**Image processing and 3D reconstruction**

Movie frames of the *Rl*GabD•NADH were motion corrected without binning, using a pixel size of 0.574 Å, and dose-weighted using the Motioncorr2 program.^1^ Contrast transfer function (CTF) corrections were performed using CTFFIND 4.1.^2^ Most of the subsequent processing steps were carried out using RELION 3.^3^ Laplacian-of Gaussian (LoG) based automated particle picking was performed on the data. Particles were extracted and subjected to 2D classification. 2D classes showing sharp structural features were chosen to build an initial 3D model. This initial model was used for template-based picking of particles (low pass-filtered at 20 Å). Picked particles were 2D classified to remove poor particles. Particles were 3D classified, without the use of symmetry constraints. The class showing well-defined structural features was then selected for 3D refinement with D2 symmetry imposed. This gave a reconstruction with a resolution of 3.37Å (FSC threshold of 0.143). Subsequently, CTF refinement^4^ was performed for magnification anisotropy; optical aberrations (up to the fourth order); and per-particle defocus and per-micrograph astigmatism. Bayesian Polishing^5^ was also used to optimise per-particle beam-induced motion tracks, followed by another round of auto-refinement. CTF refinement, Bayesian Polishing and 3D refinement steps were repeated to yield a 2.52 Å map (FSC threshold of 0.143). Further details of the image processing and 3D reconstruction can be found in Table S1 and Fig. S6.

**Model building, refinement, and validation**

An initial model was built using the map_to_model function in Phenix.^6^ The initial model was modified using *Coot*.^7^ Cycles of real space refinement in *Phenix* and interactive model building in *Coot* were used to fit the structure to the map.^6^ Model geometries were assessed with MolProbity.^8^ Structures and maps in the Fig. were rendered with ccp4mg,^9^ PyMOL (http://www.pymol.org/) or ChimeraX.^10^

**Sequence similarity network analysis**

Sequence Similarity Network (SSN) analysis was carried out using the web-based Enzyme Function Initiative Enzyme Similarity Tool (EFI-EST) (https://efi.igb.illinois.edu/efi-est/),^11^ and Genome neighborhood analysis was carried out using the web-based EFI Genome Neighborhood Tool (EFI-GNT (https://efi.igb.illinois.edu/efi-gnt/).^12^ All networks were visualized using Cytoscape v3.9.^13^ The SSN created using the alignment score 150 was used to generate genome neighbourhood diagrams (GND) with open reading frame (ORF) ± 10 neighbours using the EFI-GNT. A script was used to extract the accession codes of the retrieved neighbours. The SLA dehydrogenase neighbours were used to generate an SSN of neighbours (SSNN; **Fig. S18**). The resulting SSNN was plotted using Cytoscape v3.9, with minimum alignment score of 50. We used the SSNN to build a genome neighborhood similarity network (GNSN) in which each node corresponded to a single SLA dehydrogenase that was connected by an edge to another SLA dehydrogenase if they shared >4 isofunctional genes in their genome neighborhood. This condition results in clustering of nodes involved in similar SQ degradation pathways. To annotate the GNSN we coloured the nodes according to whether the SLA dehydrogenase gene possessed specific neighboring genes encoding SQ degradation pathway enzymes in the genome neighborhood, namely the gateway enzyme SQase (PF01055) for the sulfo-EMP (SF kinase YihV, PF00294), sulfo-EMP2 (SF kinase SqiK, PF00365), sulfo-ED (SG dehydratase, PF00920), sulfo-SFT (SF transaldolase, PF00923), and DHPS dehydrogenase (DHPS degradation pathway, PF03446), as well as manual inspection. A separate GNSN was coloured according to phylum of the host bacterium.

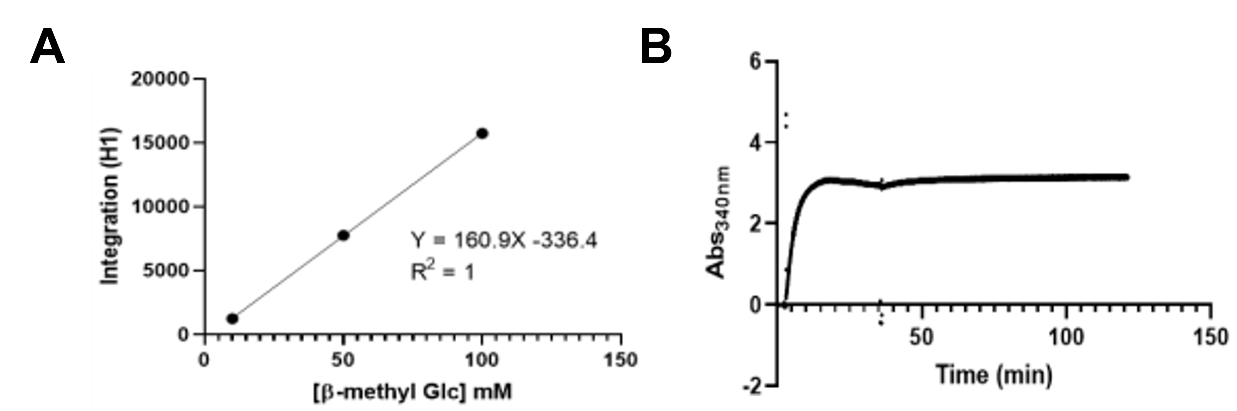

**Figure S1. A)** Calibration curve of the integration of the anomeric proton in the ^1^H NMR spectrum of methyl β-glucoside as a function of concentration, used for determination of [SLA]. **B)** Representative plot (performed in triplicate) for increase in absorption (corresponding to production of NADH) from oxidation of SLA catalyzed by *Rl*GabD SLA dehydrogenase monitored using UV-Vis spectrophotometer at 340 nm. Reactions contained 30 mM Tris buffer pH 7 at 30 °C [NAD^+^] = 1.5 mM and [SLA] = 1 mM, and were initiated by addition of *Rl*GabD to final concentration of 15.8 nM. The reaction appeared complete by 40 min. At this time, more *Rl*GabD was added to a final concentration of 205 nM, and the reaction was monitored for 100 min with no further increase in absorbance. Maximum absorbance (A_340_) = 3.081, corresponding to consumption of 48.7% of the SLA.

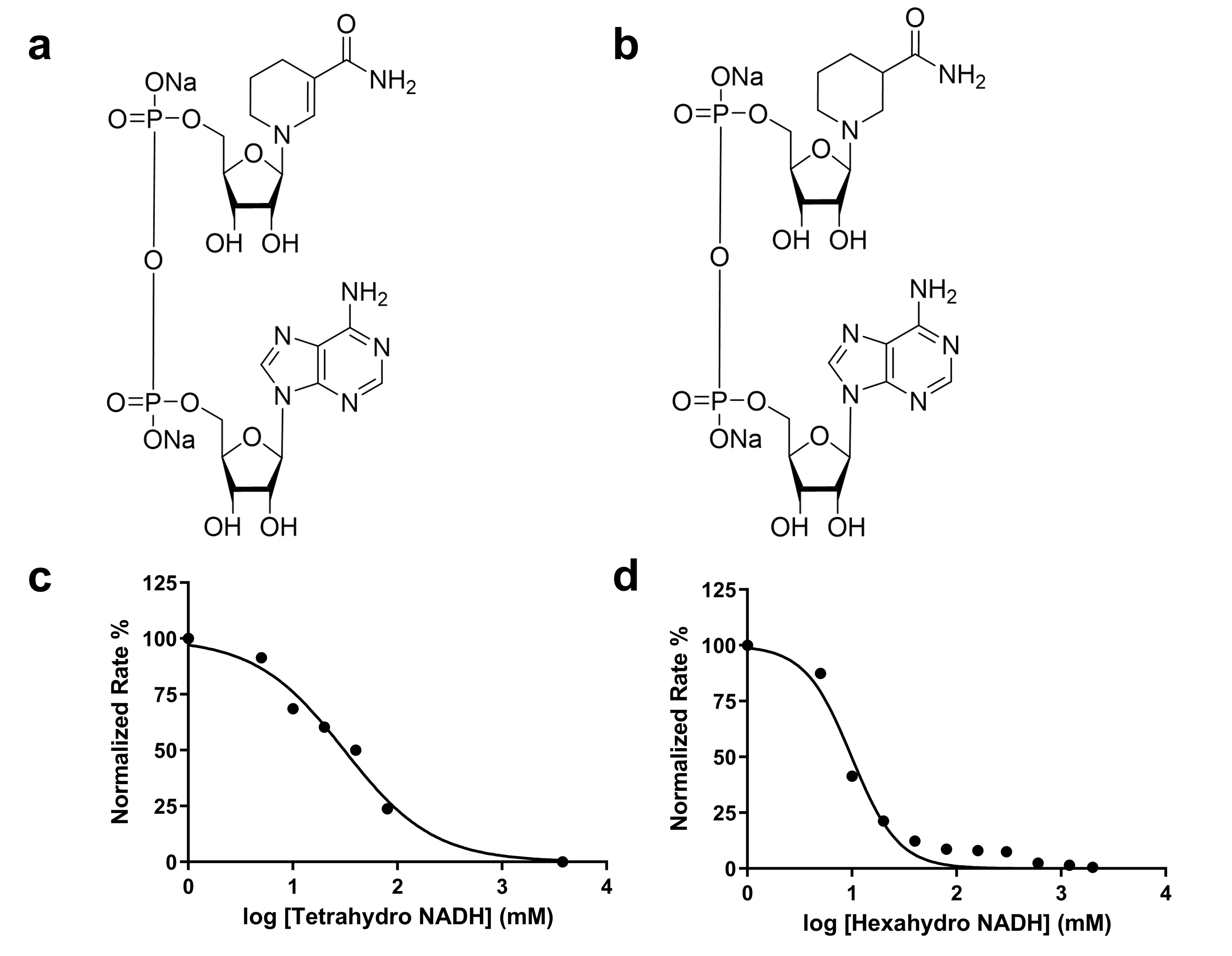

**Figure S2.** Structures of **a)** tetrahydro-NADH and **b)** hexahydro-NADH. Tetrahydro-NADH and hexahydro-NADH were assessed as inhibitors of *Rl*GabD. IC_50_ values were determined in 30 mM Tris buffer and 30 mM KCl (pH 7.12) with 0.1% BSA at 30 °C using constant [SLA] (at *K*_M_^SLA^ /10) and constant [NAD^+^] (at *K*_M_^NAD+^) using a UV/Vis spectrophotometer at 340 nm. **c)** Inhibition of *Rl*GabD at constant NAD^+^ and SLA by tetrahydro NADH, IC_50_ (tetrahydro-NADH) = 28.2 μM; and **d)** hexahydro NADH, IC_50_ (hexahydro-NADH) = 9.1 μM.

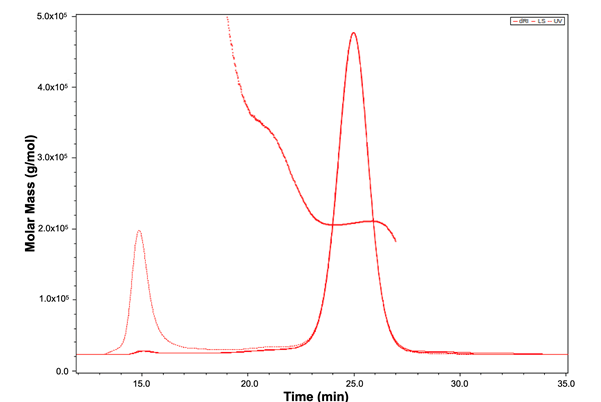

**Figure S3.** **SEC-MALLS plot reveals the oligomeric state of *Rl*GabD in solution.** UV-trace and an average molecular weight trace (red), calculated from the refractive index and light scattering signal gave mass estimation of 207 kDa, which corresponds to a homotetramer (with monomer molecular weight of 51568 Da). The area eluted under the major peak corresponds to 337 µg, which comprises ~96% of the eluted material confirming near homogeneity of the sample. A small amount of material at the void volume has MW >8 MDa, corresponding to some larger aggregates.

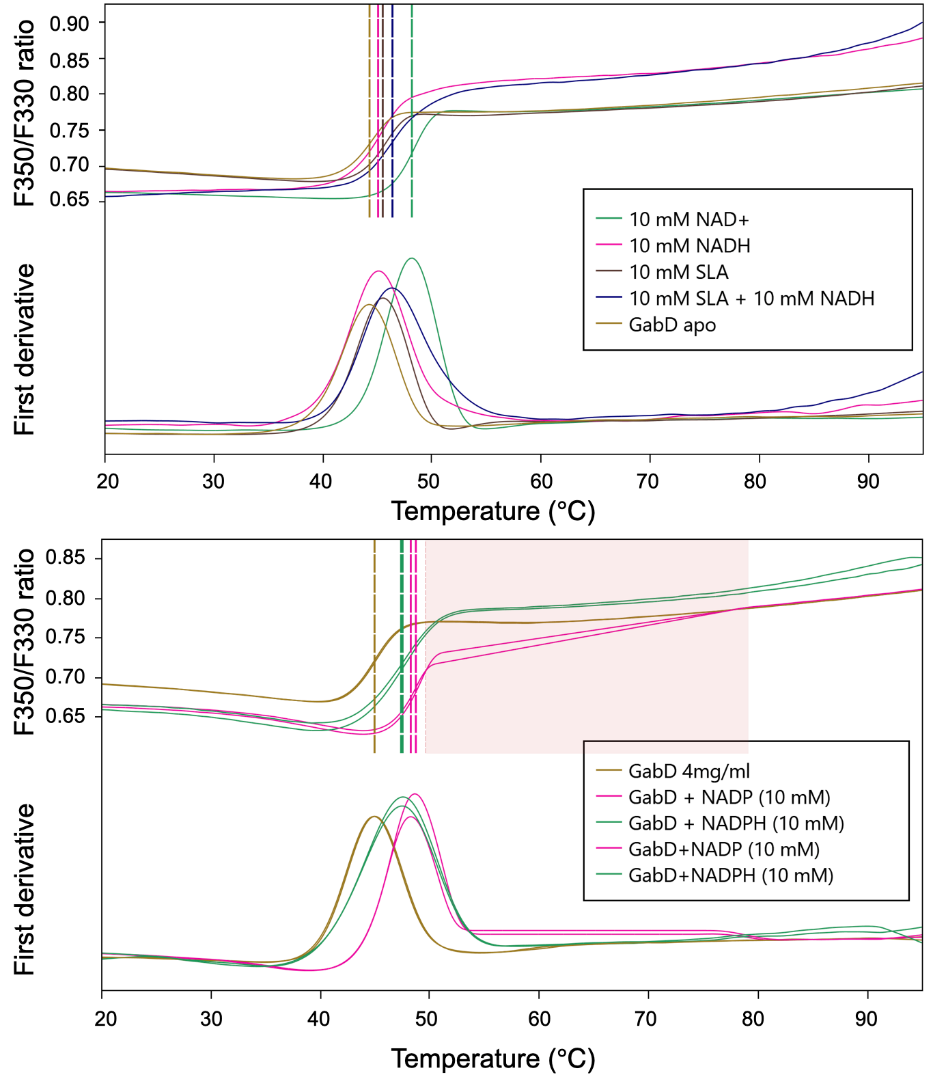

|  | **Sample** | **Inflection point for F350/F330 (Replicate 1)** | **Inflection point for F350/F330**  **(Replicate 2)** | **Trend** | **∆Tm (Avg)** |
| --- | --- | --- | --- | --- | --- |
|  | *Interaction of RlGabD with substrates SLA, NAD(H) and analogs* | | | | |
| 1 | RlGabD apo 4mg/mL | 44.2°C | 44.0°C |  |  |
| 2 | RlGabD + NAD (10 mM) | 48.1°C | 48.8°C | ↑ | 4.3°C |
| 3 | RlGabD + NADH (10 mM) | 45.1°C | 45.6°C | ↑ | 1.5°C |
| 4 | RlGabD + SLA (10 mM) | 45.5°C | 45.4°C | ↑ | 1.3°C |
| 5 | RlGabD + NADH + SLA (10 mM each) | 46.4°C | 47.1°C | ↑ | 2.6°C |
| 6 | RlGabD + NADH3 (10 mM) | 43.7°C | 43.9°C | ↓ | 0.3°C |
| 7 | RlGabD + NADH4 (10 mM) | 43.6°C | 43.9°C | ↓ | 0.4°C |
|  | *Interaction of RlGabD with NADP(H)* |  |  |  |  |
| 1 | RlGabD apo 4mg/ml | 45.0°C | 44.9°C |  |  |
| 2 | RlGabD + NADP (10 mM) | 48.7°C | 48.3°C | ↑ | 3.5°C |
| 3 | RlGabD + NADPH (10 mM) | 47.5°C | 47.4°C | ↑ | 2.5°C |

**Figure S4.** **NanoDSF studies with *Rl*GabD.** Tm shifts occurring upon binding cofactors NAD(P)H (and analogs) and substrate SLA.

B.

A.

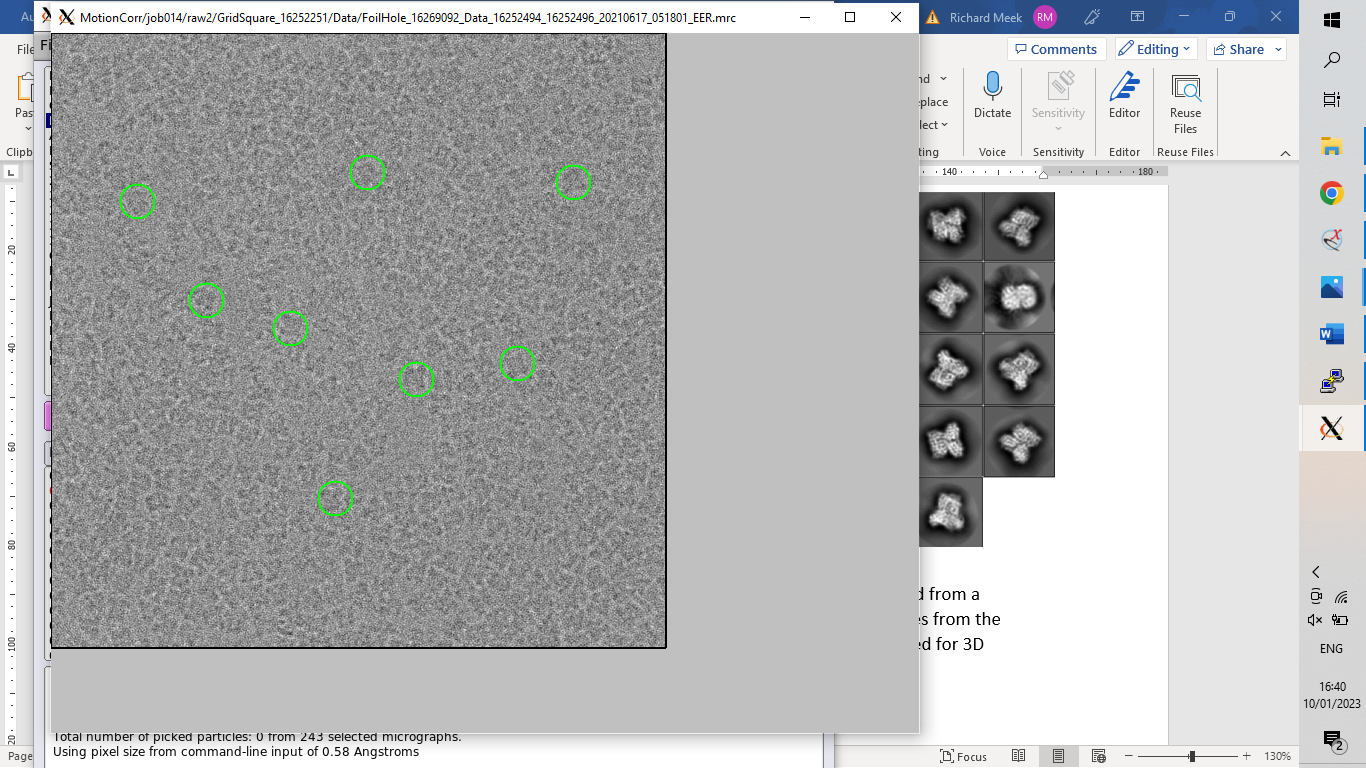

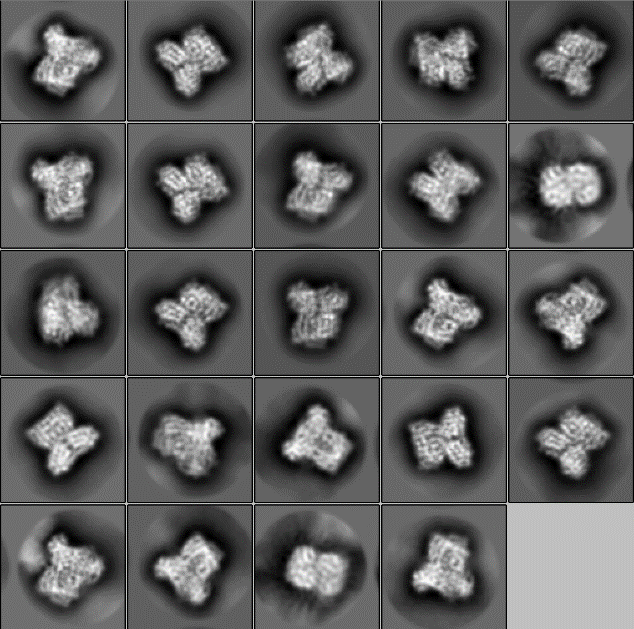

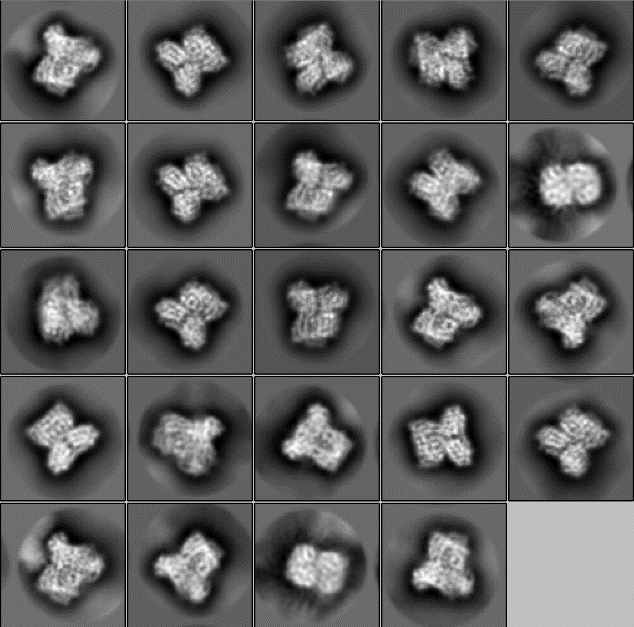

**Figure S5. Cryo-EM data processing.** (A) Representative micrograph of GabD collected from an UltrAuFoil R1.2/1.3, 300 mesh gold grid. Green circles highlight a few picked particles from the micrograph. White scale bar is 150 Å. Total micrographs collected = 934. (B) 2D class averages of GabD used for 3D classification.

Raw micrographs (934)

Motion correction

CTF estimation

**A1**

Micrographs (865)

LoG-based Autopicking

Particles (396276)

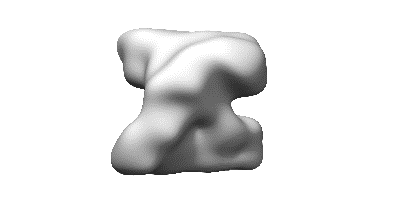

2D Classification

Initial model

Low-pass filter: 20 Å

Particles (302938)

Template Autopicking

Particles (437107)

2D Classification

Particles (269200)

3D Classification

75165

66677

60492

**Particles**

66866

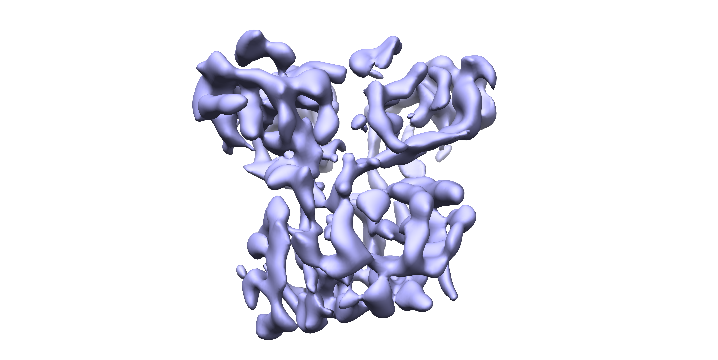

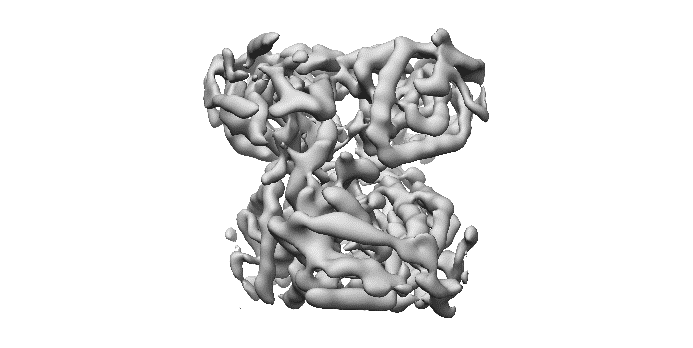

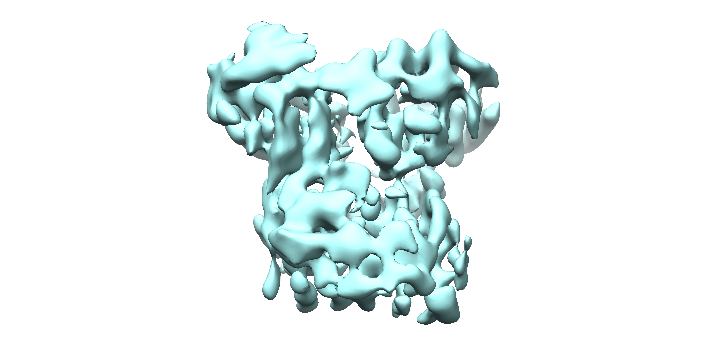

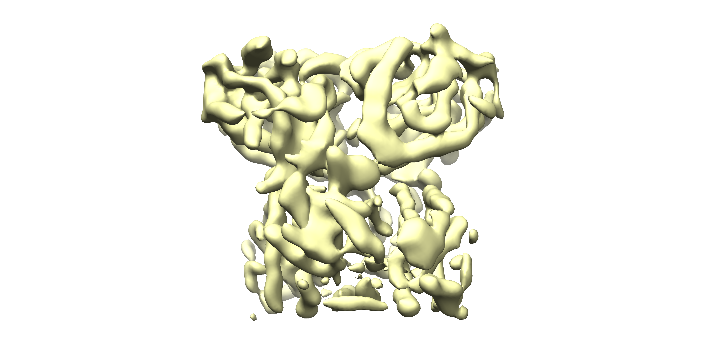

2.45 Å

4.26 Å

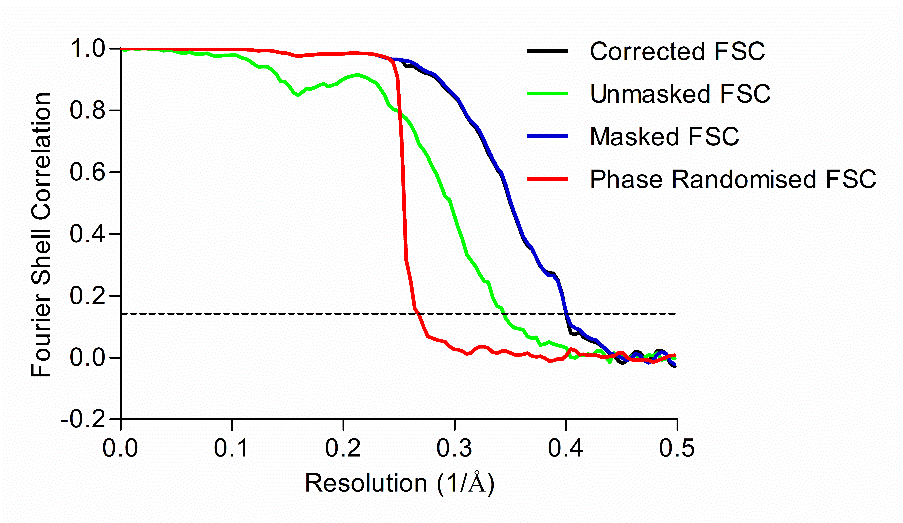

**B1**

CTF refinement

Bayesian Polishing

3D Refinement

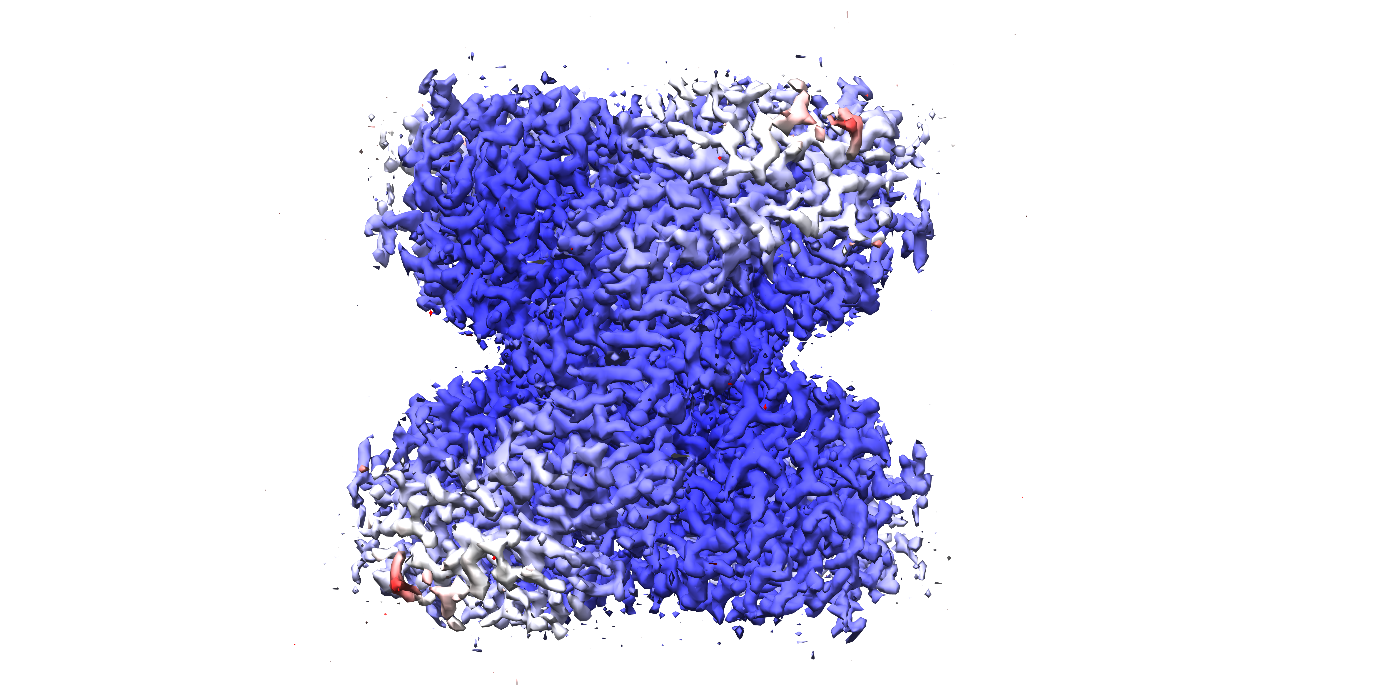

**C1**

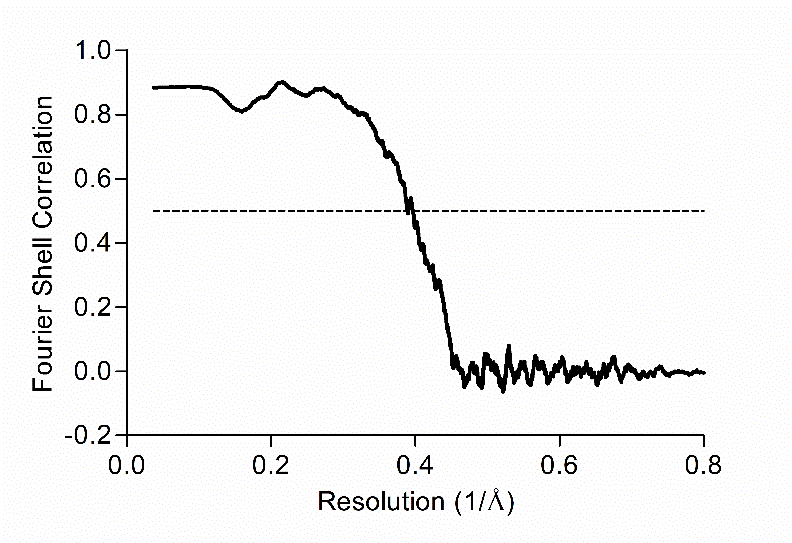

**Figure S6.** **Data processing and refinement workflow of *Rl*GabD Cryo-EM structure.** (**A**) Data processing steps from initial data collection to final reconstruction. Numbers in Parentheses refer to total number of micrographs or particles. Final reconstruction coloured by local resolution as estimated through Relion. (**B**) Fourier Shell Correlation (FSC) between the two independently refined maps. Horizontal dotted black line, FSC=0.143 (**C**) FSC between the final map reconstruction and *Rl*GabD model. Horizontal dotted black line, FSC=0.5.

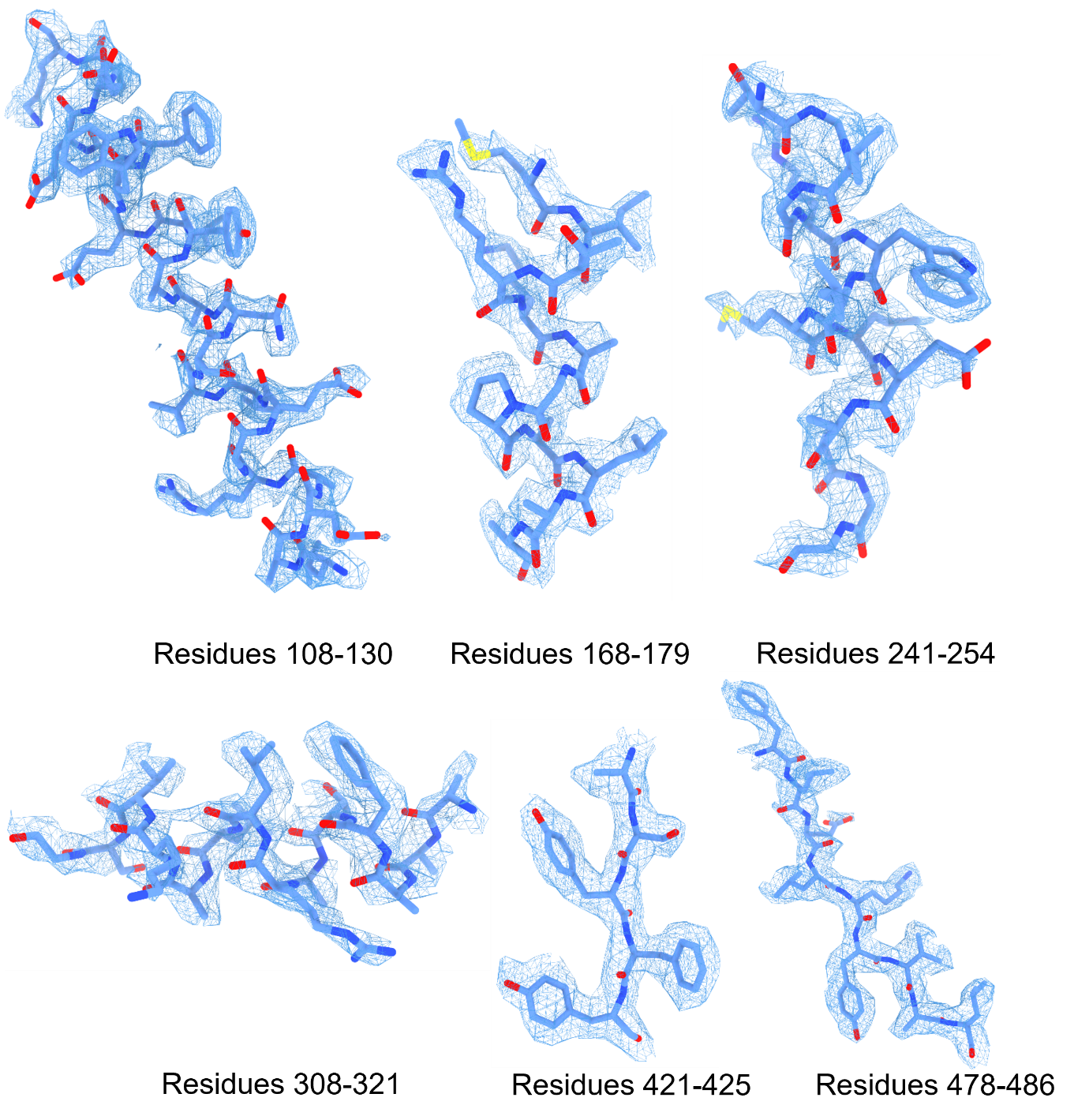

**Figure S7.** **Representative electron densities of different regions of *Rl*GabD.** Densities carved at a 1.4-1.6 Å radius around residues at a threshold of 0.04-0.05.

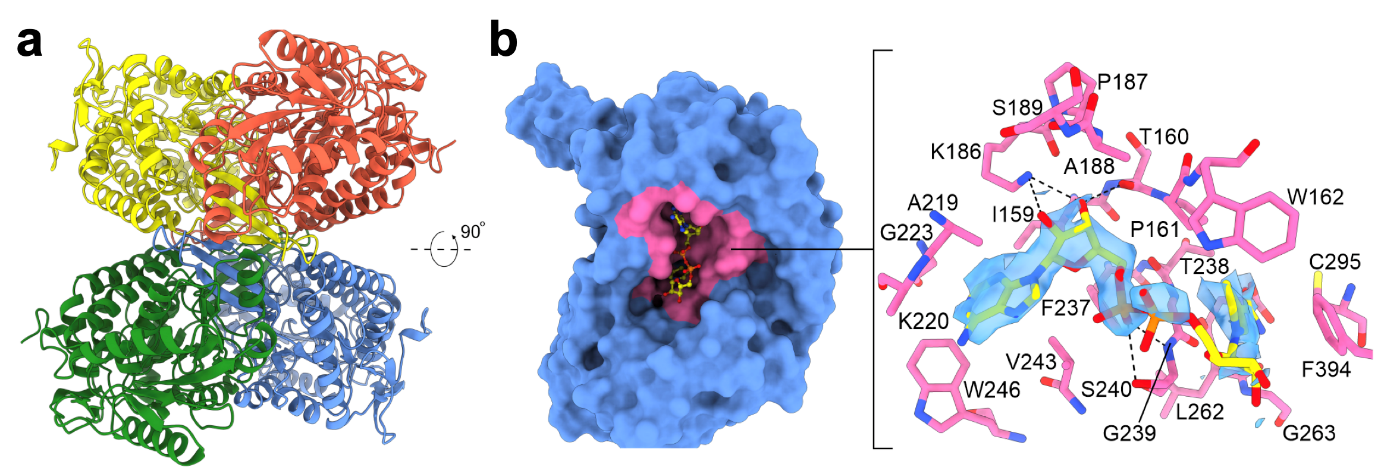

**Figure S8. Cryo-EM structure of *Rl*GabD•NADH complex.** a) Quaternary structure of *Rl*GabD•NADH complex shown in cartoon representation. b) Surface representation of *Rl*GabD•NADH complex showing the NADH binding site (left); residues lining the NADH binding site are shown in stick format. Blue mesh is the density map at a threshold of 0.037.

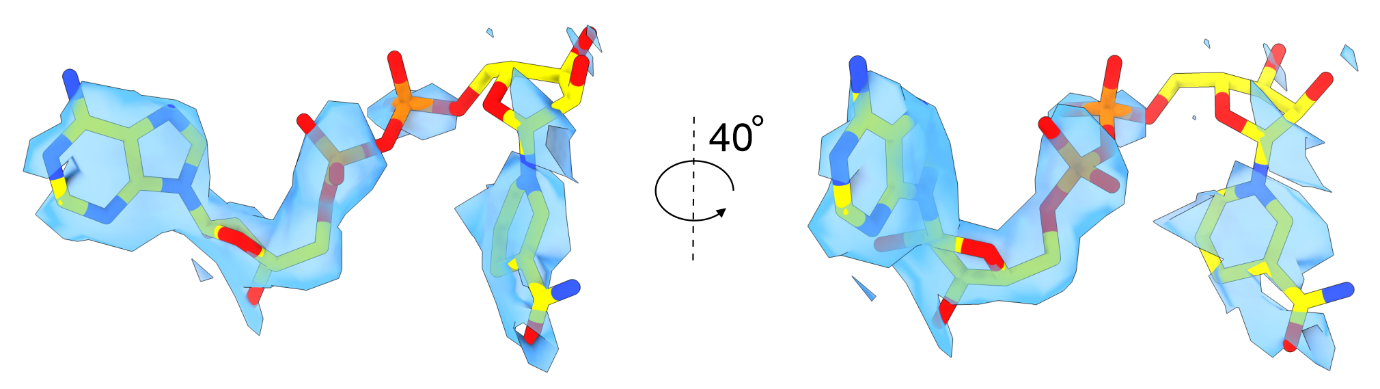

**Figure S9. Ligand density for NADH bound to *Rl*GabD.** While the density is well defined for the adenosine group and attached phosphate, the density is poor for the second phosphate and nicotinamide group. The density is almost totally absent for the phosphate-to-nicotinamide linking ribose.

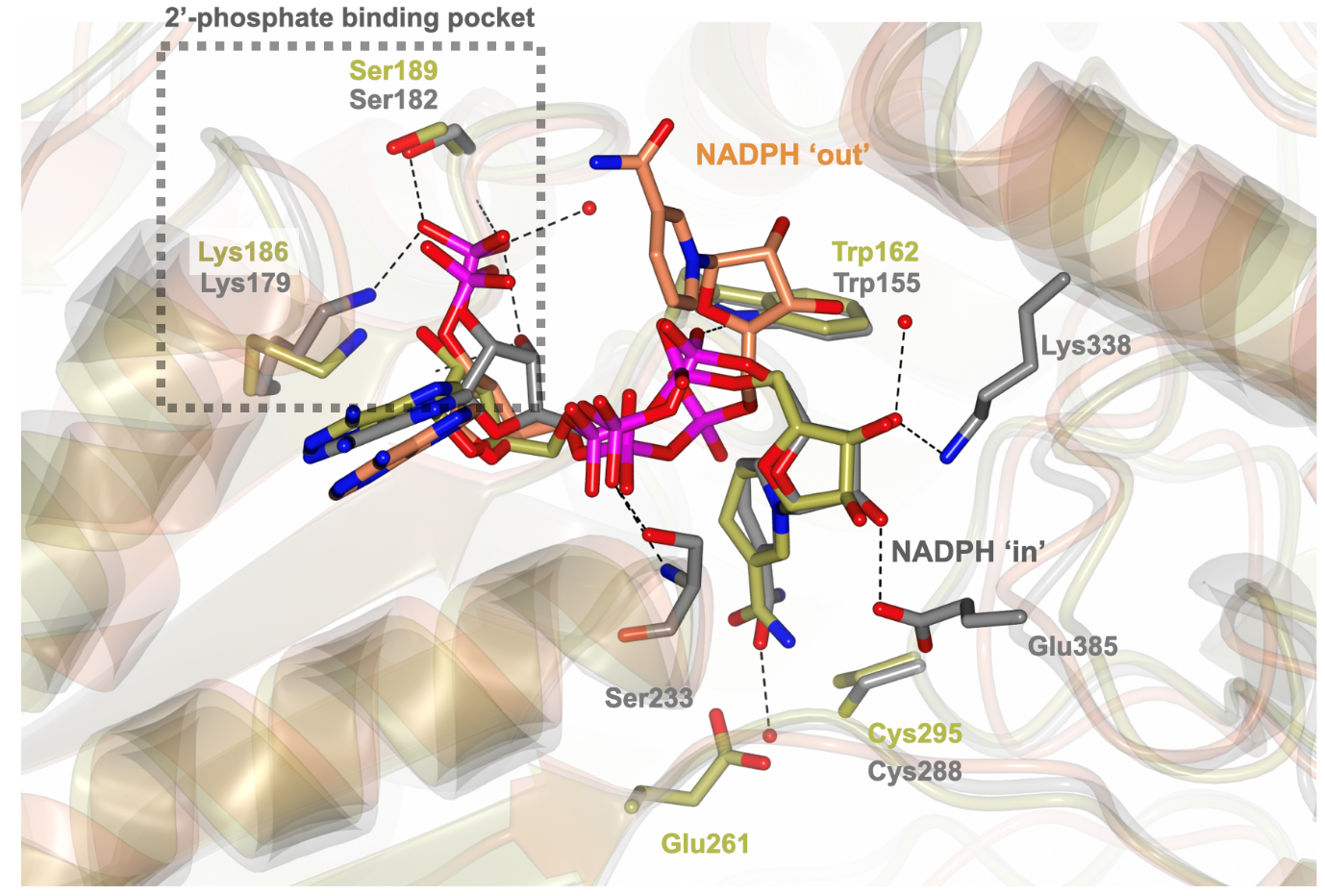

**Figure S10.** Superposition of crystal structures of binary complexes of *Rl*GabD•NADH (in gold), *E. coli* SSADH•NADPH (PDB: 3JZ4) in grey and *E. coli* lactaldehyde dehydrogenase•NADPH (PDB: 2ILU) in coral showing the 2’-phosphate binding pocket and different conformations of the nicotinamide ring of NAD(P)H cofactor in aldehyde dehydrogenases.

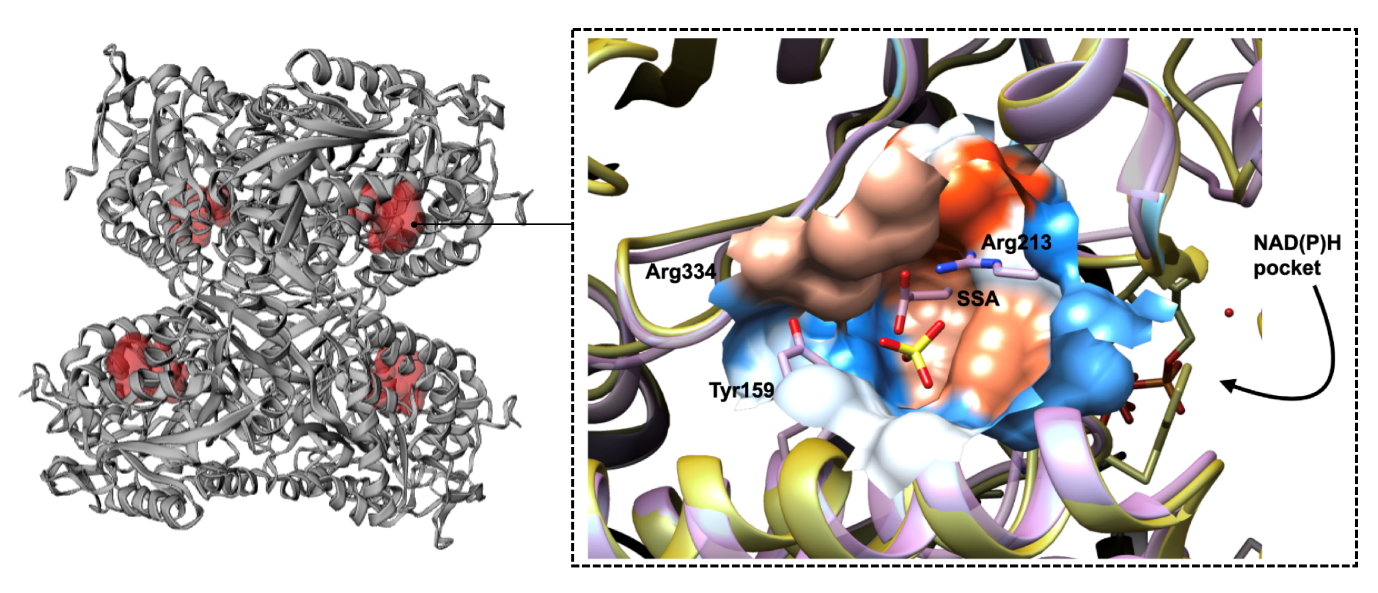

**Figure S11.** Imprint of sulfonate substrate binding pockets depicted in *Rl*GabD tetramer (in grey, left), as computed by CastP 3.0 server shown in red. Overlay of *Rl*GabD substrate (SLA) binding pocket with the crystal structure of human SSADH with bound succinic semialdehyde (SSA) (pdb: 2w8q). *Rl*GabD and human SSADH are shown in yellow and purple, respectively, and residues in the vicinity of bound SSA are shown in purple.

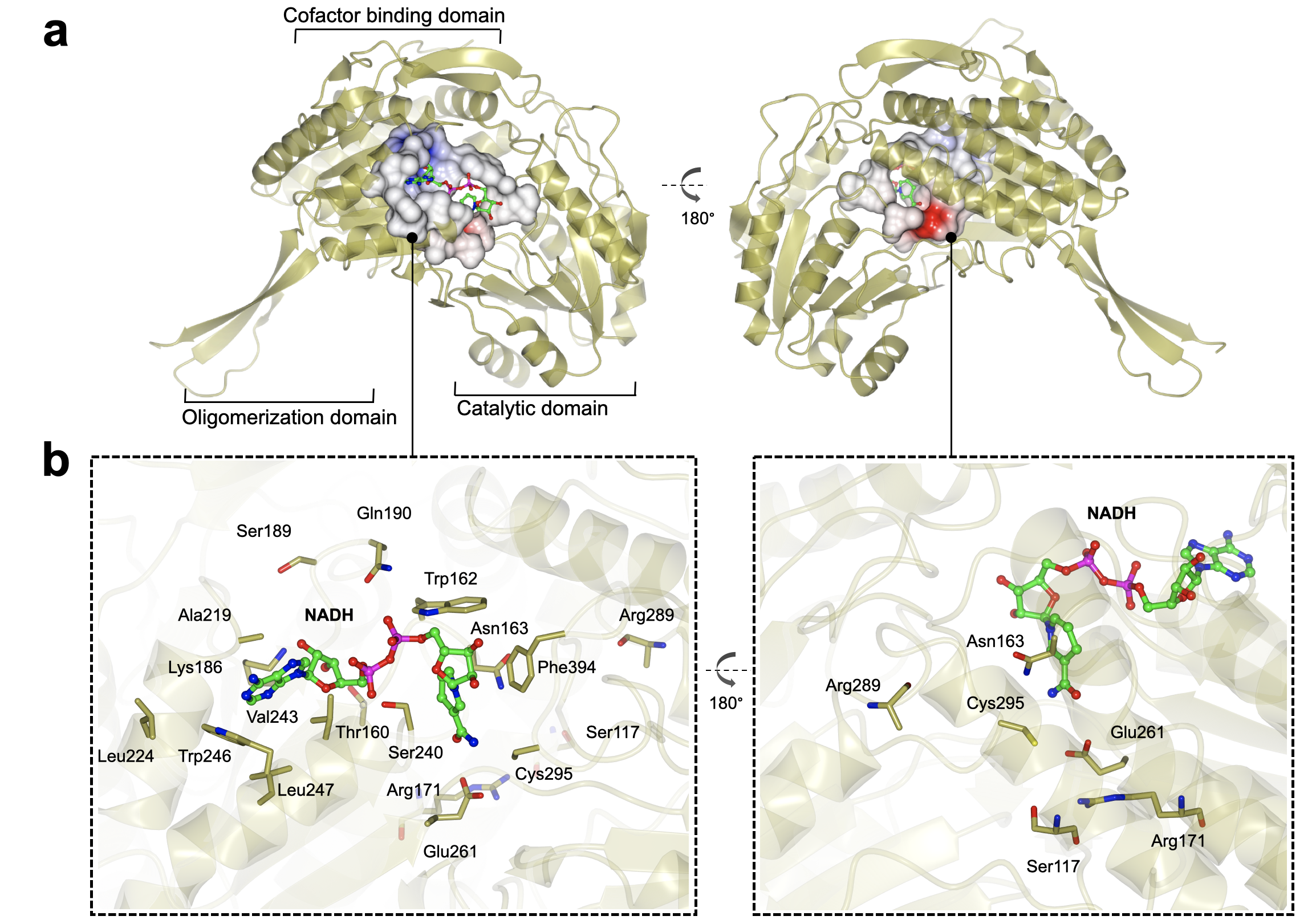

**Figure S12. Additional views of NADH binding pocket from front and back. a)** *Rl*GabD subunit showing the three-domain architecture and the two views of the NADH binding pocket represented as the electrostatic potential of the residues lining the pocket. The red region corresponds to the catalytic Glu-Cys dyad. **b)** Close up view of the NAD(H) binding pocket depicting the residues lining the NAD(H) and SLA binding pockets.

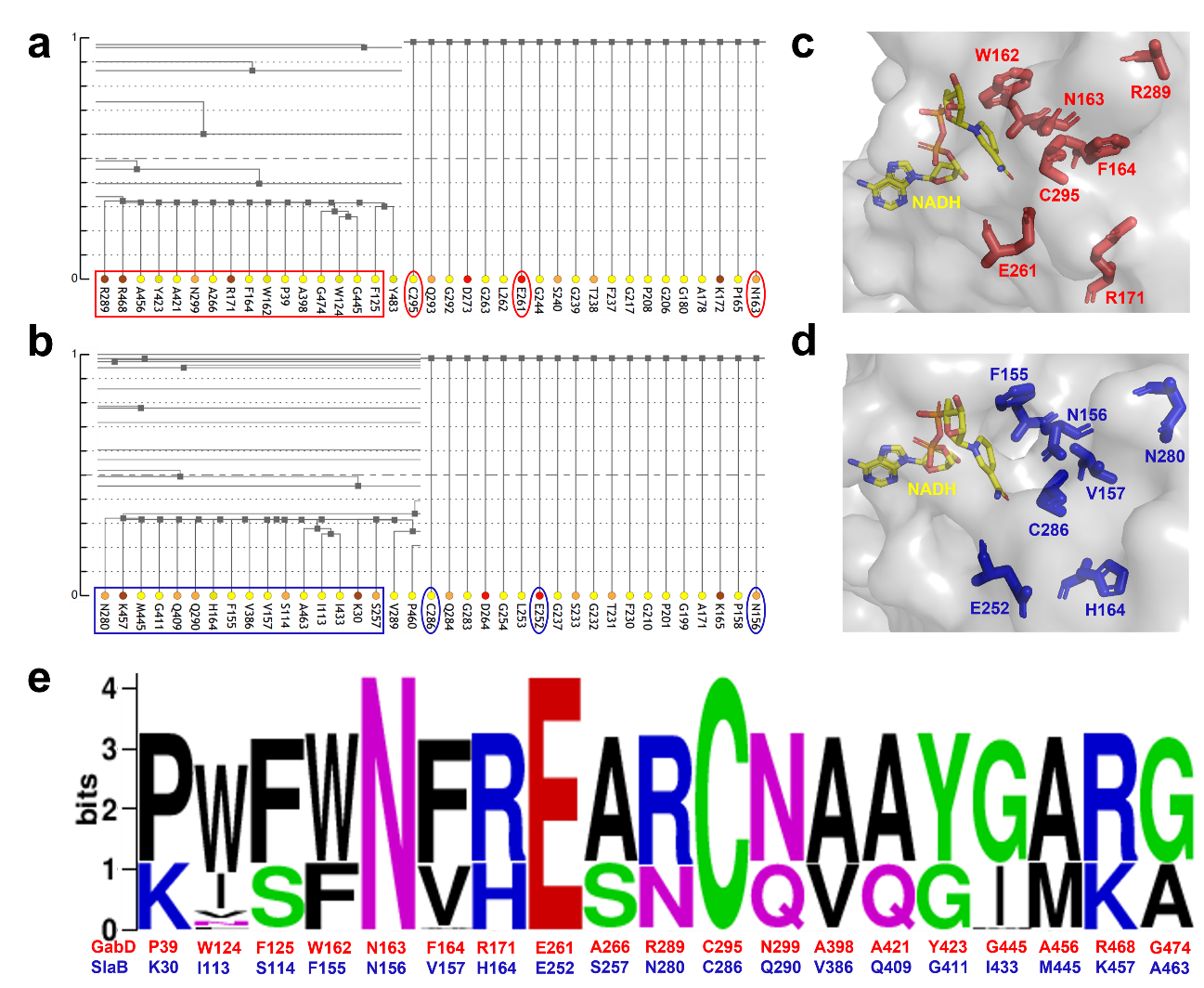

**Figure S13.** Co-evolution analysis of SLADH. Snapshot of cluster tree diagrams of **a)** *Rhizobium leguminosarum* GabD and **b)** *Bacillus megaterium* SlaB, generated by CoeViz.^14^ The cluster trees were generated using the complete linkage hierarchical clustering algorithm based on Mutual Information (MI), with scores are converted to distances by 1-score transformation.^15^ Circled residues are conserved residues involved in the reaction mechanism. Boxed residues are identified as co-evolving with R171/R289 (*Rl*GabD) and H164/N280 (*Bm*SlaB). Location of conserved mechanism residues, and co-evolved residues proximal to the proposed SLA binding site shown on the **c)** cryo-EM structures of *Rhizobium leguminosarum* GabD and **d)** the AlphaFold2 predicted structure of *Bacillus megaterium* SlaB. **e)** Sequence logo of conserved mechanism residues and residues co-evolving with R171/R289 (*Rl*GabD) and H164/N280 (*Bm*SlaB), generated by WebLogo (https://weblogo.berkeley.edu/logo.cgi)

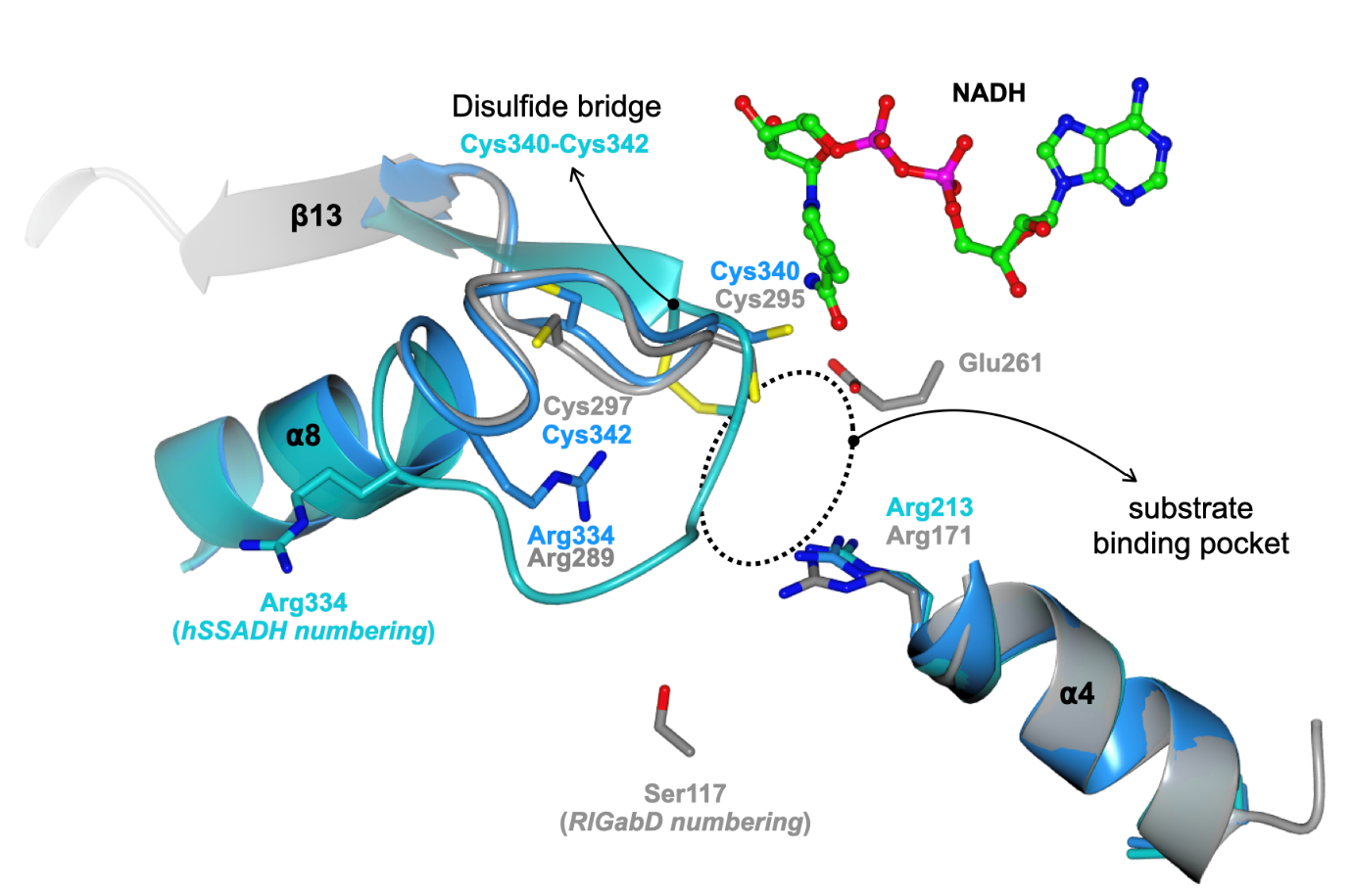

**Figure S14.** Overlay and close-up view of active site of SLA dehydrogenase *Rl*GabD compared to human SSADH in oxidized ‘closed’ (PDB: 2W8N, shown in turquoise) and reduced ‘open’ (PDB: 2W8O, shown in blue) forms. In hSSADH the loop containing the catalytic cysteine nucleophile can undergo redox modulation accompanied with large loop position changes due to reversible disulfide bridge formation with adjacent cysteine. In GabD, the equivalent loop is present in the ‘open’ form with active site residues poised for substrate binding and catalysis. NB: in our structure of *Rl*GabD the side-chain of Arg289 was disordered and was not modelled.

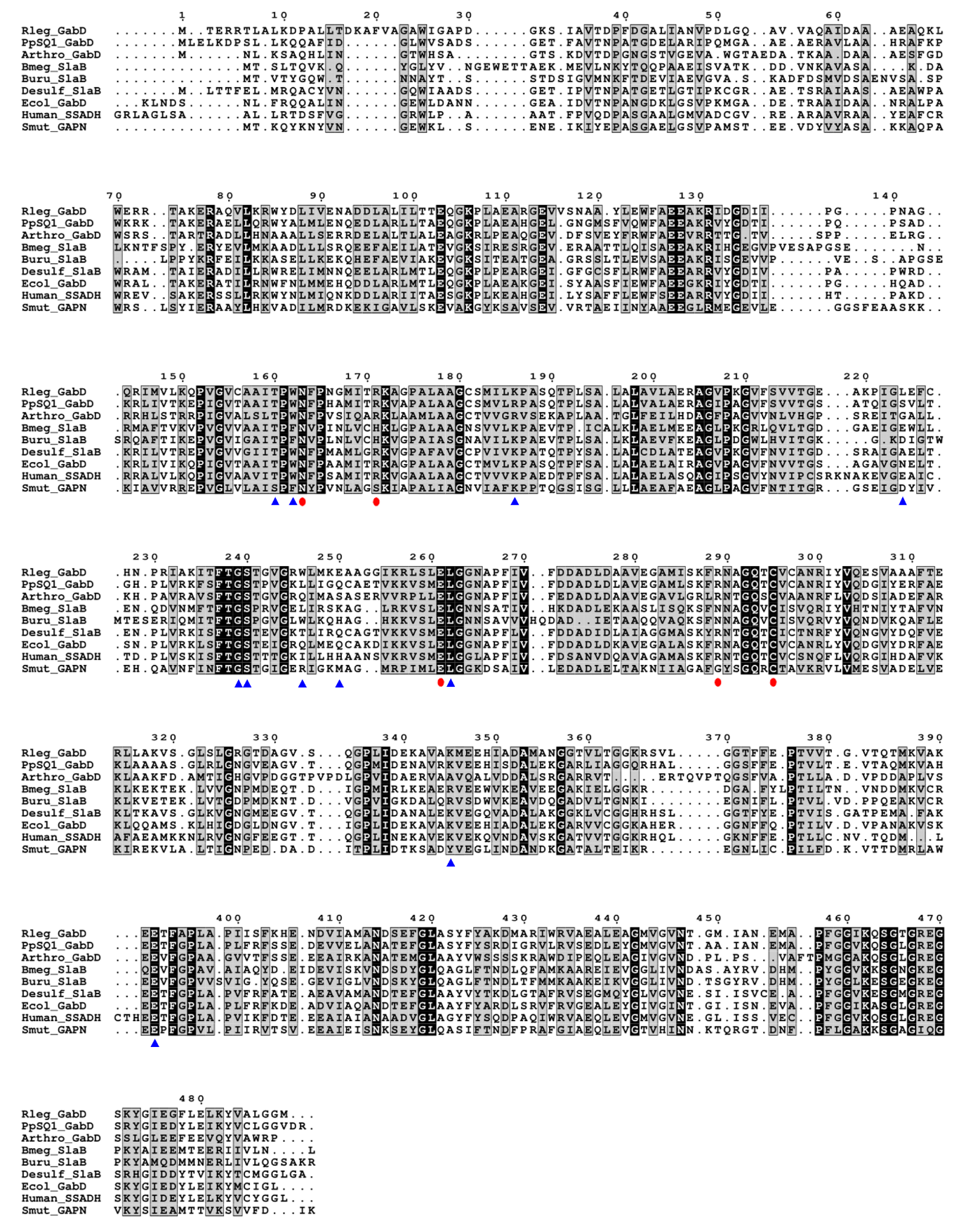

**Figure S15.** Multiple sequences alignment of SlaB/GabD proteins including SSADH (*E. coli*) and SLA dehydrogenases (*Arthrobacter* sp.*, Rhizobium leguminosarum*, *Pseudomonas putida*, *Herbaspirillum seropedicae*, *Bacillus megaterium*, *Bacillus urumqiensis)*. Secondary structural elements are annotated from the structure of *E. coli* SSADH (PDB 3JZ4). Annotated by red dots are residues shown to be important in the catalytic mechanism. The residues of NADP^+^ binding to *E. coli* SSADH are annotated by blue arrows.

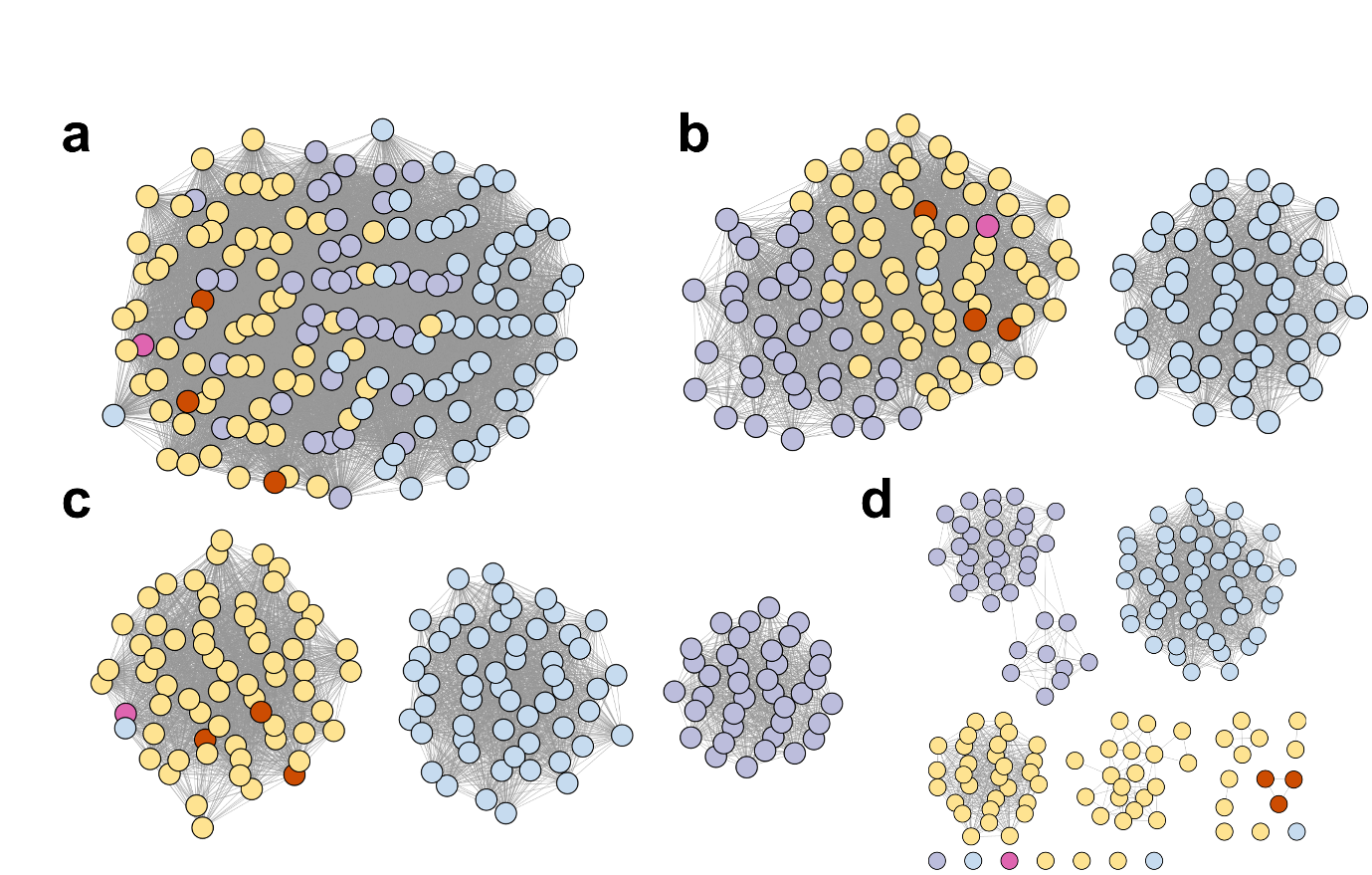

**Figure S16.** SSNs created using different alignment scores. **a)** At alignment score 75, the sequences form a single cluster. b) At alignment score 100, the sequences form two clusters. c) At alignment score 125-150, the SSN breaks into three clusters that separate into three main Phyla, with Chloroflexi and Candidatus Dormibacteraeota segregating with Actinobacteria. d) At higher alignment threshold (175), the SSN produces a large number of singletons. Colouring scheme: Actinobacteria (yellow), Chloroflexi (red), Candidatus Dormibacteraeota (pink), Firmicutes (light blue), Proteobacteria (light purple).

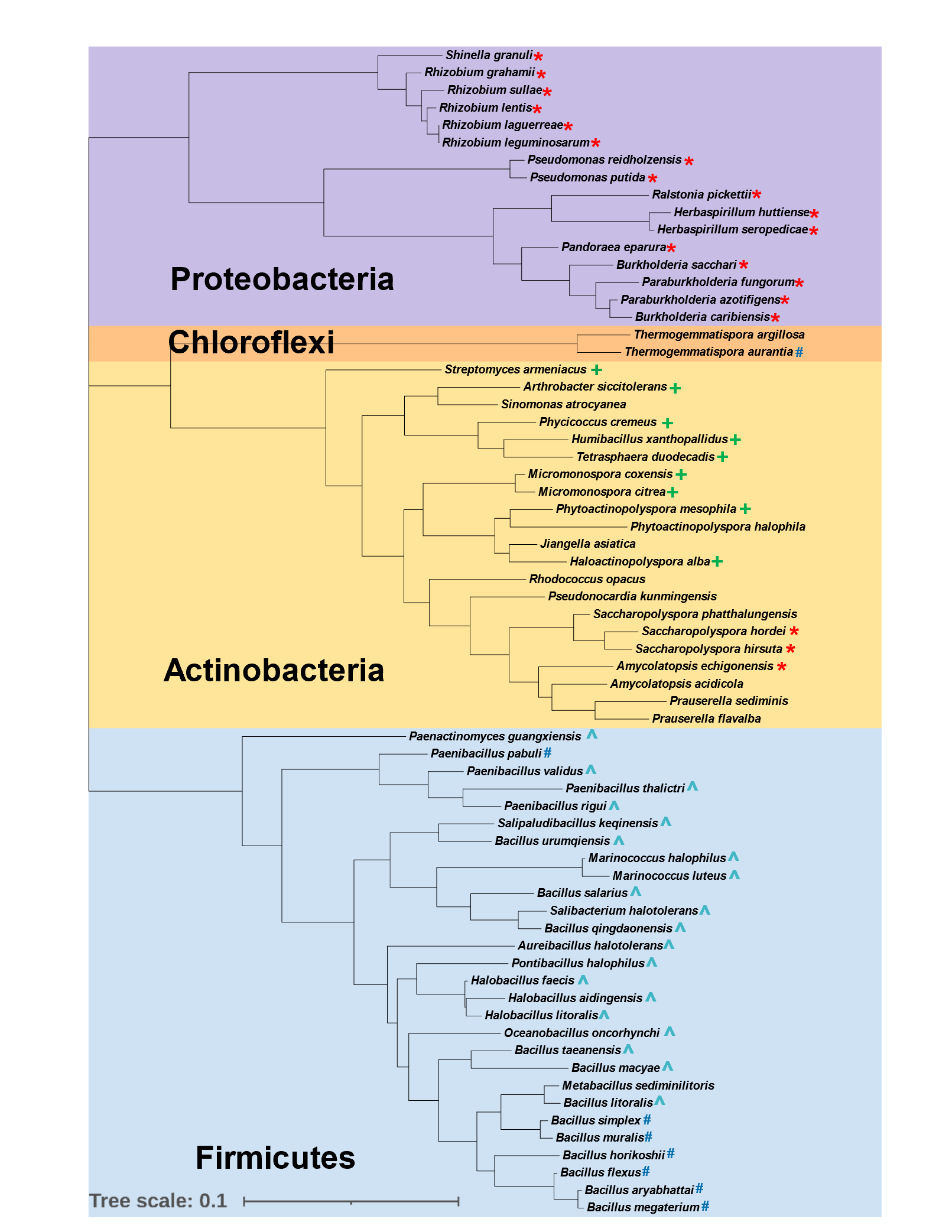

**Figure S17.** A phylogenetic tree based on 16S ribosomal RNA of organisms from four different phyla containing putative SLADH proteins in 4 different sulfoglycolytic pathways: sulfo-ED (*****), sulfo-SFT (**#**), sulfo-EMP2 (**^**) and sulfo-EMP3 (**+**). Note: Candidatus could not be included as its 16S ribosomal RNA data was unavailable.

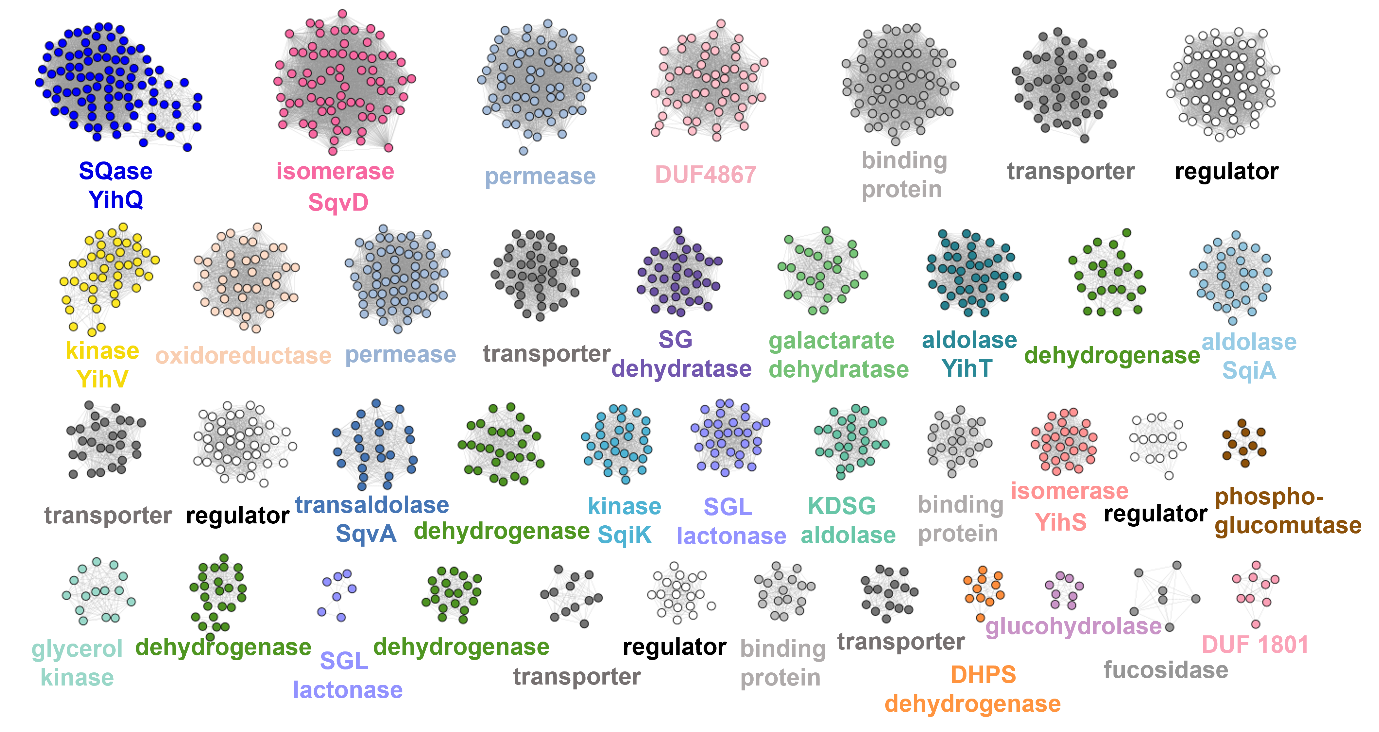

**Figure S18.** SSN of SLADH neighbors, displayed at alignment score of 50 while the percent identity is 40 and the length of the sequences are between 45 to 1151. Clusters are colored according to their functions. Key proteins from the four sulfoglycolytic pathways are: SQ isomerase (YihS, pink; SqvD, red pink), SQase (blue), aldolase (SqiA, light blue; KDSG aldolase, light green), SF kinase (YihV, yellow PF00294; SqiK, aquamarine PF00365), dehydratase (SG dehydratase, purple PF00920), lactonase (SGL lactonase, light purple), transaldolase (SqvA, dark blue PF00923).

**Table S1.** Data collection, processing and refinement statistics for *Rl*GabD•NADH complex structure

|  | *Rl*GabD complexed with NADH (PDB:8C54) |
| --- | --- |
| Data collection and processing | |
| Nominal magnification | 240,000 |
| Voltage (kV) | 200 |
| Electron exposure (e-/Å^2^) | 50 |
| Defocus range (μm) | -2, -1.75, -1.5, -1.25, -1.0, -0.9, -0.7 |
| Pixel Size (Å) | 0.574 |
| Symmetry imposed | D2 |
| Initial number of particles | 437,107 |
| Final number of particles | 75,165 |
| Map resolution (Å) (FSC threshold) | 2.52 (0.143) |
| Refinement | |
| Initial model used (PDB) | N/A |
| Model resolution (Å) (FSC threshold) | 2.57 (0.5) |
| Map sharpening *B* factor (Å^2^) | -44.85 |
| *Model composition* |  |
| Non-hydrogen atoms | 14266 |
| Protein residues | 1928 |
| Ligands | 4 |
| *RMSDs* |  |
| Bond lengths (Å) | 0.003 |
| Bond angles (°) | 0.488 |
| *Ramachandran plot* |  |
| Outliers (%) | 0 |
| Allowed (%) | 2.2 |
| Favoured (%) | 97.8 |
| *Validation*  Rotamer outliers (%) | 3.43 |
| MolProbity | 1.82 |
| Clash score | 6.47 |
| Model vs. Data (CC mask) | 0.84 |
